# Dynamic modelling of EWS::FLI1 fluctuations reveals molecular determinants of phenotypic tumor plasticity and prognosis in Ewing sarcoma

**DOI:** 10.1101/2025.04.03.647002

**Authors:** Veveeyan Suresh, Christoph Hafemeister, Andri Konstantinou, Sarah Grissenberger, Caterina Sturtzel, Florencia Cidre-Aranaz, Andrea Wenninger-Weinzierl, Martha Magdalena Zylka, Karla Queiroz, Dorota Kurek, Ana Sastre, Javier Alonso, Martin Distel, Anna Obenauf, Thomas G.P Grünewald, Florian Halbritter, Heinrich Kovar, Valerie Fock

## Abstract

The mechanisms underlying tumor cell plasticity driving drug resistance and disease progression remain poorly understood. In Ewing sarcoma (EwS), variations in EWS::FLI1 (EF) activity have been associated with epithelial-mesenchymal plasticity (EMP). Using degron technology, we titrated endogenous EF in an EwS cell line and linked phenotypic states to distinct EF thresholds. Strikingly, modest EF depletion promoted a pro-metastatic phenotype, that diminished upon near-complete EF loss. Nascent RNA sequencing revealed distinct gene clusters with heterogenous response patterns to varying EF dosage. Target genes most sensitive to subtle EF depletion contained GGAA microsatellites in EF-bound enhancers. Furthermore, we identified Krüppel-like zinc-finger transcription factors associated with EF-repressed EMP genes. Transient EF depletion followed by rapid restoration to simulate oncoprotein fluctuations identified persistently dysregulated genes associated with poor prognosis. This study underscores the therapeutic challenge of insufficient EF inhibition and provides a foundation for exploiting oncoprotein dynamics to uncover therapeutic vulnerabilities in fusion-driven cancers. Beyond EwS, our results underscore the broader impact of oncoprotein dosage dynamics in cancers with otherwise quiet genomes.

**SIGNIFICANCE:** We report EwS as a paradigm for the importance of oncogene fluctuations in tumor cell plasticity and disease progression. Effective therapeutic strategies must ensure complete EF depletion to prevent inadvertent metastasis.

## INTRODUCTION

Ewing sarcoma (EwS) is an aggressive bone and soft tissue cancer predominantly affecting adolescents. Despite intensive multimodal therapy, the five-year survival rate for patients with relapsed or metastatic disease remains only 20-30% (1). This highlights the urgent need for the development of more effective treatment strategies. Among potential therapeutic targets, the EWS::FLI1 fusion protein (EF) expressed in 85% of EwS cases has emerged (2,3). EF functions as an aberrant ETS family transcription factor that binds to consensus sites containing a GGAA core motif in promoters and enhancers, and exhibits dual activity as a transcriptional activator and repressor (4,5). By rewiring the epigenome, EF can turn bound GGAA microsatellites into *de-novo* enhancers activating nearby genes in an EwS-specific manner (5–7). EF-driven gene repression occurs either directly through interactions with co-repressors, or indirectly by activating other transcriptional repressors and recruiting repressive epigenetic complexes (8–11). This leads to the downregulation of genes critical for mesenchymal identity and affects tumor cell plasticity (12,13).

EwS is characterized by quiet genomes with few recurrent mutations (14–17) but exhibits a significant degree of epigenetic plasticity and intra-tumoral heterogeneity (18). Importantly, the existence of multiple tumor cell phenotypes in EwS has been linked to variations in the levels and activity of EF (19–21). For instance, single-cell RNA-profiling studies of EwS patient-derived cell lines, xenografts, and immunohistochemistry on primary tumors identified a subpopulation of ∼1-2% dormant-like cells which were associated with low EF expression (19,21). EF knockdown models suggest that high EF levels are linked to a proliferative phenotype, whereas cells with low EF expression display mesenchymal features and an increased propensity for migration and metastasis (19,22–25). EF activity is modified by loss of STAG2 which frequently occurs on a subclonal level and is associated with adverse prognosis (26,27). EF activity is also subject to competition with other transcription factors such as ETV6 and HOXD13 at shared chromatin binding sites (28–30). Additionally, regulation on the transcriptional and RNA/protein stability levels directly affect EF expression (31–38).

Fluctuations in EF levels may occur stochastically or in response to microenvironmental cues (18). At the cell population level, these fluctuations can lead to diverse phenotypic states and intratumoral heterogeneity (39,40). Factors such as hypoxia, serum starvation, and signaling through Rho or Hippo pathways can impact EF expression and activity (19,22,23). However, a comprehensive model that captures the full spectrum of EF expression and activity levels in EwS has been lacking. This knowledge gap hinders the development of novel therapeutic strategies aimed at overcoming the inherent metabolic and metastatic plasticity of EwS cells.

In this study, we resolved how distinct thresholds in EF protein levels influence gene regulation and functional cell states in EwS. We engineered an EwS cell line to enable precise, tunable and reversible modulation of EF dosage using the dTAG system (41). This allowed us to establish a gradient of distinct EF thresholds to describe the short-term and long-term phenotypic and molecular consequences of EF fluctuations at defined amplitudes. We identified a set of persistently dysregulated genes involved in EMP, which were associated with poor patient prognosis in a cohort of 196 EwS patients. Together, these results highlight the crucial role of EF fluctuations in EwS cell plasticity and disease progression. They underscore the importance of completely depleting EF in EwS tumors as insufficient or transient inhibition may unintentionally promote the development of lethal metastases.

## RESULTS

### Degron knock-in for tunable EF expression from the endogenous gene locus

To precisely modulate endogenous EF dosage in EwS cells, we made use of the dTAG system which allows rapid and dose dependentprotein degradation (41). We employed a CRISPR genome editing approach to tag endogenous EF in A673 EwS cells with the fluorescent protein mNeonGreen (mNG), serving as a quantitative proxy for EF levels. Additionally, we introduced an FKBP12^F36V^ tag, facilitating targeted degradation of the fusion protein upon treatment with the heterobifunctional von Hippel-Lindau (VHL)-binding degrader molecule, dTAG^V^-1 (Fig. 1A). Using FACS-mediated single-cell sorting followed by clonal expansion, we obtained clones with biallelic knock-in of the mNG-HA-FKBP12^F36V^ tag at the C-terminal end of the *FLI1* coding region (EF-dTAG clones) (Supplementary Fig. S1A). Targeting the EF locus in other EwS cell lines (TC71, STA-ET-7.2) resulted in duplication of the EF allele with only one copy successfully modified, indicating selective pressure against this EF manipulation (data not shown). Focusing on A673 EF-dTAG clones A2.2 and B3.1, western blot analysis showed that the absolute levels of tagged EF were similar between the clones and comparable to their untagged wildtype counterpart (Supplementary Fig. S1B). Consistent with the absence of expression of the unrearranged *FLI1* allele, no signal was obtained for wildtype FLI1 for both the parental cells and EF-dTAG clones (Supplementary Fig. S1B). CUT&RUN revealed similar genome-wide chromatin binding patterns between untagged and tagged EF in parental cells and A673 EF-dTAG clone A2.2, respectively (Supplementary Fig. S1C, Supplementary Table S1). RNA-seq analysis identified 564 and 302 differentially expressed genes (DEGs) in EF-dTAG clones A2.2 and B3.1, compared to parental A673 cells, respectively, with an enrichment of genes linked to epithelial-mesenchymal transition (EMT) (Supplementary Fig. S1D, Supplementary Table S2). A similar number of DEGs was detected when comparing individual untagged A673 single-cell clones with the parental bulk population, suggesting that transcriptional differences arise from clonal heterogeneity or the single-cell cloning process (Supplementary Table S2). The EF-dTAG clones exhibited moderately reduced proliferation (Supplementary Fig. S1E), accompanied by an increase in migration and invasion, compared to the parental A673 cells (Supplementary Fig. S1F). The A2.2 dTAG clone exhibited colony-forming ability in soft agar comparable to that of the parental cells, whereas the B3.1 cells showed a reduced capacity for anchorage-independent growth (Supplementary Fig. S1G).

**Figure 1.**
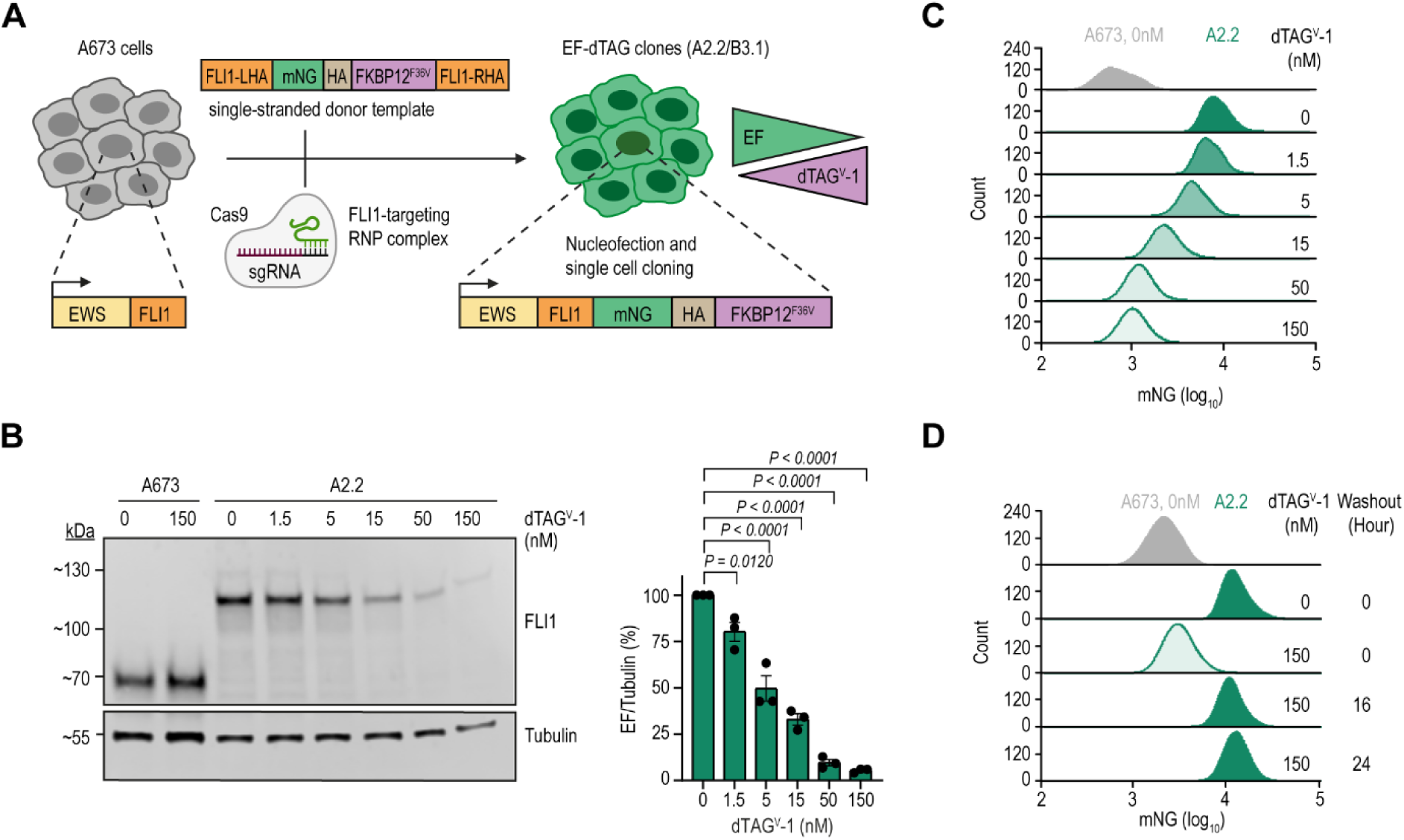
Generation and establishment of EF thresholds. **A**, Schematic representation of the A673 cell line editing strategy and EF titration using the dTAG system. sgRNA, single-guide RNA; RNP, ribonucleoprotein; mNG, mNeonGreen. **B**, Western blot analysis of FLI1 protein levels in EF-dTAG clone A2.2 and parental A673 cells treated with the indicated concentrations of dTAG^V^-1 for 24 hours (left). Tubulin was used as a loading control. Bar graphs depict EF protein levels normalized to Tubulin, presented as mean ± SEM (right; *n* = 3 independent experiments). *P*-values were determined using one-way ANOVA with post-hoc Dunnett’s multiple comparisons test. **C**,**D** Flow cytometry analysis of mNG fluorescence intensity in EF-dTAG clone A2.2 treated with increasing concentrations of dTAG^V^-1 for 24 hours (**c**) or following 16 or 24 hours of washout after treatment with 150 nM dTAG^V^-1 (**D**). Data shown are representative of at least three independent experiments.

To titrate EF levels, we treated EF-dTAG clones with a 3-fold dilution series of dTAG^V^-1 concentrations in the nanomolar range and assessed EF protein levels by immuno blot, flow cytometry, and confocal microscopy (Fig. 1B-C and Supplementary Fig. S2A-C). We observed a dTAG^V^-1 concentration-dependent gradual and highly reproducible loss of EF protein, decreasing by approximately 14% (1.5 nM), 35% (5 nM), 57% (15 nM), 84% (50 nM) and 93% (150 nM) at 24 hours in both EF-dTAG clones (Fig. 1B, Supplementary Fig. S2A,C and Table 1). Upon ligand washout, EF levels returned to baseline levels within 16 hours (Fig. 1D). Together, these results validate our engineered dTAG clones as a suitable model for studying EF fluctuations in EwS cells.

**Table 1.** Quantification of EF protein levels following dTAG^V^-1 treatment for 24 hours.

| Mean EF protein values (n=3) as quantified by Western blot (WB) or confocal microscopy using mNeonGreen as proxy(CF) |  |  |  |  |  |  |  |  |  |  |  |  |
| --- | --- | --- | --- | --- | --- | --- | --- | --- | --- | --- | --- | --- |
| dTAG<sup>V</sup>-1 conc (nM) | 0 |  | 1.5 |  | 5 |  | 15 |  | 50 |  | 150 |  |
| Clone/Methods | WB | CF | WB | CF | WB | CF | WB | CF | WB | CF | WB | CF |
| A2.2 | 100.000 | 100.000 | 80.213 | 91.667 | 49.570 | 72.333 | 32.919 | 50.667 | 9.617 | 9.666 | 5.431 | 0.333 |
| B3.1 | 100.000 | 100.000 | 87.983 | 86.000 | 63.948 | 72.666 | 38.027 | 52.333 | 16.637 | 27.000 | 5.370 | 15.000 |
| **Average of methods** |  |  |  |  |  |  |  |  |  |  |  |  |
| Clone/dTAG<sup>V</sup>-1 conc (nM) | 0 | 1.5 | 5 | 15 | 50 | 150 |  |  |  |  |  |  |
| A2.2 | 100 | 86 | 61 | 42 | 10 | 3 |  |  |  |  |  |  |
| B3.1 | 100 | 87 | 68 | 45 | 22 | 10 |  |  |  |  |  |  |
| Consolidated % EF left | 100 | 86 | 65 | 43 | 16 | 7 |  |  |  |  |  |  |
| % EF reduction | 0 | 14 | 35 | 57 | 84 | 93 |  |  |  |  |  |  |

### Partial EF depletion significantly increases migration and invasion with loss of colony formation *in vitro*

To investigate the impact of distinct EF levels on functional phenotypes in EwS, we pre-treated EF-dTAG clones with increasing concentrations of dTAG^V^-1 ligand for 24 hours, followed by Transwell migration assays (Fig. 2A). We found that a stepwise reduction of EF protein levels resulted in a gradual increase in the number of migratory EwS cells peaking at 15 nM dTAG^V^-1 (≈ 43% EF, Table 1). Further EF depletion began to reverse the effect on cell migration. Consistently, the invasive potential of dTAG clones into collagen I, assessed using an organ-on-a-chip system, increased progressively with stepwise EF depletion, displaying a striking vessel-like sprouting pattern of invasive EwS cells (Fig. 2B). In contrast, cell proliferation and apoptosis rates remained unchanged by increasing dTAG^V^-1 concentrations (Supplementary Fig. S3A-B). Next, we evaluated the anchorage-independent growth capacity of dTAG^V^-1treated EF-dTAG cells in soft-agar over a 3-week period as an *in vitro* surrogate of tumorigenic potential (Fig 2C). Even a slight reduction in EF levels at 1.5 nM dTAG^V^-1 (≈ 86% EF) was sufficient to decrease colony formation, while further EF depletion completely abolished it. Collectively, these results demonstrate that a moderate, but incomplete reduction of EF levels induces significant migratory and invasive characteristics in EwS cells, while anchorage-independent tumor cell growth is highly sensitive to minor decreases in EF thresholds.

**Figure 2.**
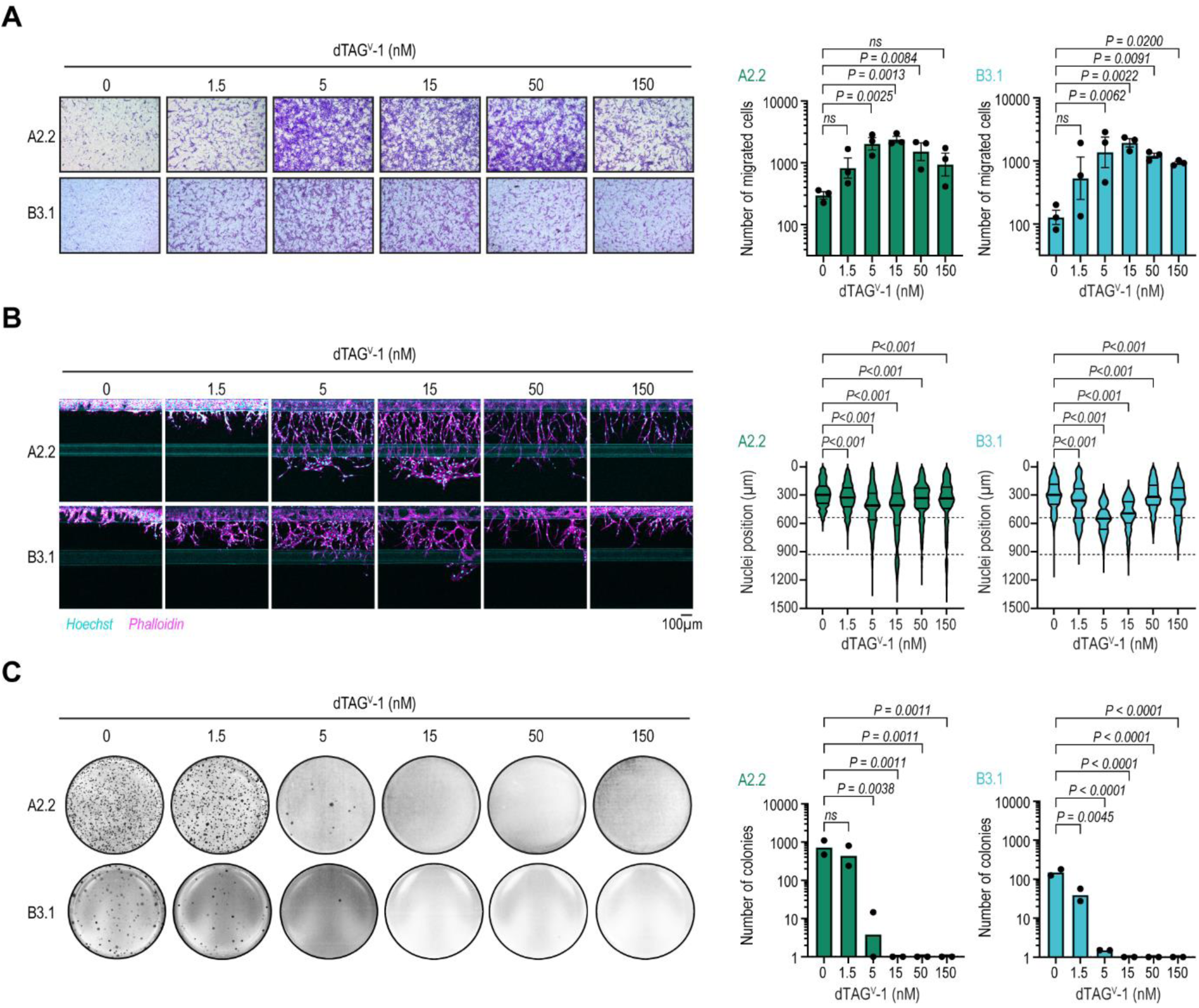
Functional characterization of the EF gradient in EF-dTAG clones. **A**, Transwell migration assay showing migrated EF-dTAG clones pre-treated with dTAG^V^-1 for 24 hours at the indicated concentrations. Representative images from one of three independent experiments are shown (left). Migrated cell counts are presented as mean ± SEM after log10 transformation (right; *n* = 3). **B**, Organoplate invasion assays depicting cell invasion into collagen I extracellular matrix (ECM) stained with Hoechst (cyan, nuclei) and phalloidin (magenta, actin cytoskeleton) 7 days after seeding and treating EF-dTAG clones at the indicated concentrations of dTAG^V^-1. Representative images from one of two independent experiments are shown (left). Nuclei positions along the Y-axis are shown, with phase guides indicated by dotted lines at 535 µm and 930 µm (right; *n* = 2). **C**, Soft agar colony formation assays showing crystal violet stained colonies formed by A2.2 and B3.1 cells three weeks after seeding and treatment with dTAG^V^-1 at indicated concentrations (Left). Representative images from one of two independent experiments are shown. Colony counts are presented as mean (*n* = 2) following log10+1 transformation. For all panels, *P*-values were calculated using one-way ANOVA with Dunnett’s post-hoc multiple comparisons test. *ns*, not significant.

### Gradual EF depletion reveals EF threshold-dependent gene sets

To explore EF level-dependent changes in the EwS transcriptome, we performed RNA-sequencing of EF-dTAG clones A2.2 and B3.1 after 24 hours of continuous dTAG^V^-1 treatment (Fig. 3A, Supplementary Table S3). A consistent transcriptomic response to increasing dTAG^V^-1 concentrations was observed for the two clones, with 1,791 genes being differentially expressed at any dTAG^V^-1 concentration in any of the two clones (DESeq2; 0nM treatment as reference; FDR < 0.05 and absolute log_2_ fold change > 2). 1,109 genes showed a gradual increase in expression, whereas 682 genes decreased upon stepwise EF depletion (Supplementary Fig. S4A-B). In line with our phenotypic observations, gene set enrichment analysis of the upregulated genes showed enrichment of gene sets related to EMT, myogenesis, hypoxia and TNF alpha signaling starting at 5 nM of dTAG^V^-1 (Fig. 3B). Conversely, E2F targets and G2M checkpoint gene sets were enriched among the downregulated genes at intermediate to high dTAG^V^-1 concentrations (>=15 nM).

**Figure 3:**
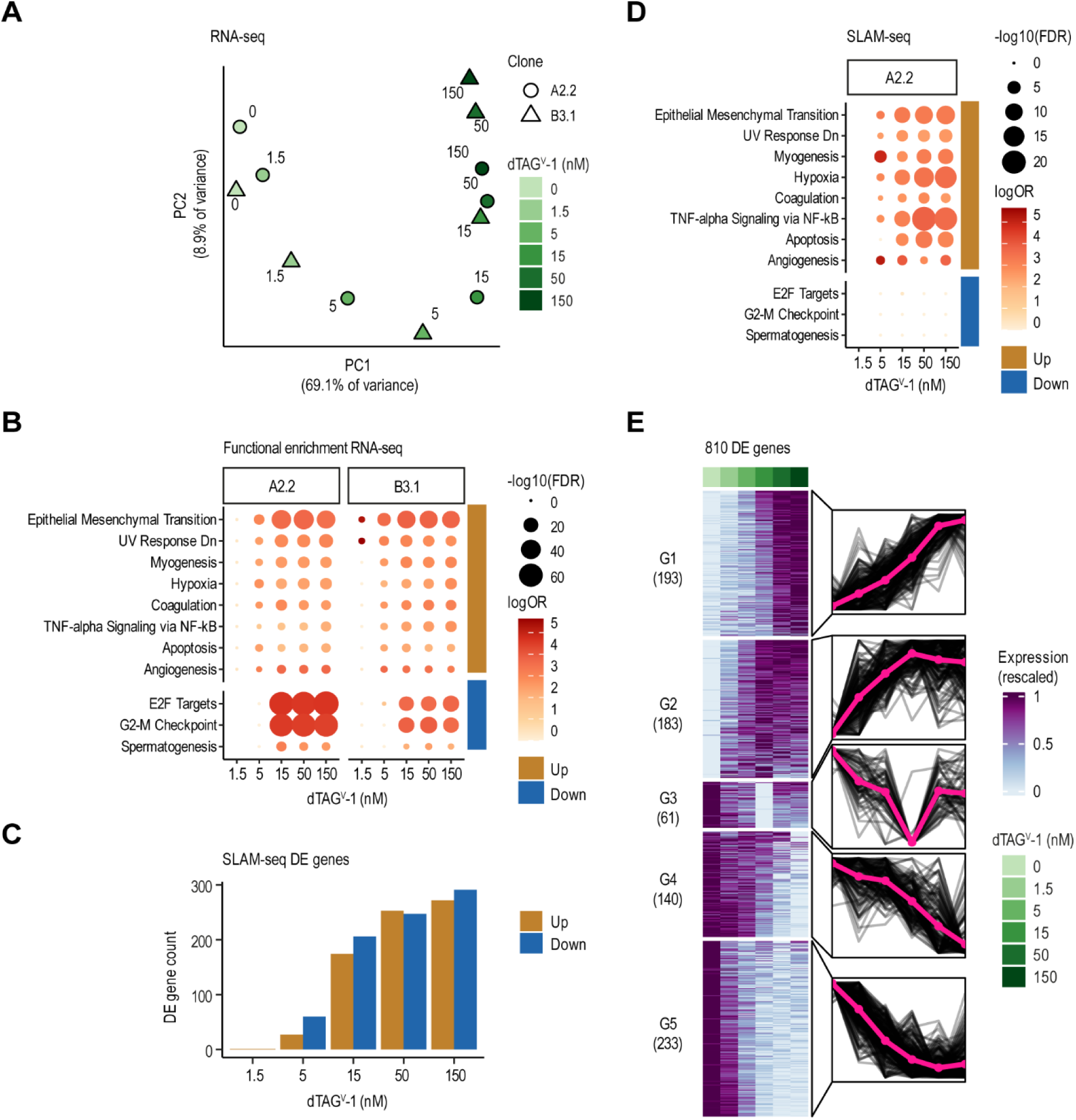
Transcriptomic alterations induced by EF gradients. **A**, EF level-dependent transcriptome changes analyzed by RNA-seq following 24-hour dTAG^V^-1 gradient treatment. Principal components of 1,791 differentially expressed genes (DEGs) across both EF-dTAG clones and dTAG^V^-1 concentrations (DESeq2^74^; 0nM treatment as reference; FDR < 0.05 and absolute log_2_ fold change > 2). Each point indicates one condition after averaging experimental replicates. Shape and color indicate the clonal cell line and the dTAG^V^-1 concentration, respectively. **B**, Functional enrichment of DEGs from panel (**A**) highlighting significantly enriched pathways from the MSigDB Hallmark 2020 database (hypergeometric test using hypeR; FDR < 0.001, log_2_ odds ratio > 3; background: all DESeq2 tested genes). Circle size and color indicate the FDR-adjusted p-value and the odds ratio, respectively. Genes were grouped by expression change direction as indicated by colored bar on the ride side. **C**, Barplots showing the number of DEGs after 3 hours of dTAG^V^-1 treatment assessed by SLAM-seq (DESeq2; FDR < 0.05, absolute log_2_ fold change > 1). Only nascent RNA reads with at least two T>C conversions were used for the analysis. **D**, Functional enrichment of DEGs from panel (**C**). Analysis as in panel (**B**). **E**, Identification of gene response groups by k-means clustering of expression profiles. Every gene from panel (**C**) was averaged across replicates, rescaled to range from 0 to 1 and plotted in a heatmap with treatment conditions on the x-axis and genes on the y-axis (left) and as individual lines grouped by cluster with conditions on the x-axis and rescaled expression on the y-axis (right). The bold pink line shows the cluster mean. The optimal number of clusters was determined using gap statistics.

To discriminate direct and indirect effects of graded EF depletion on *de novo* gene expression, we treated clone A2.2 with increasing dTAG^V^-1 concentrations for only 3 hours (Supplementary Fig. S4C) and performed thiol-(SH)-linked alkylation for the metabolic sequencing of RNA (SLAM-seq). This method enables the precise quantification of nascent RNA following dTAG^V^-1 treatment by incorporating 4-thiouridine (4SU) into transcripts resulting in thymidine-to-cytosine (T>C) conversions in RNA-sequencing reads (42,43) (Supplementary Fig. S4D, Supplementary Table S4). We found a total of 376 upregulated and 434 downregulated genes (Fig. 3C, DESeq2; FDR < 0.05, absolute log_2_ fold change > 1). Gene set enrichment analysis of the upregulated genes was consistent with the bulk RNA-seq data, revealing enrichment in pathways such as EMT, hypoxia and TNF alpha signaling (Fig. 3D). In contrast, the downregulated genes did not result in any significant enrichment.

To identify genes with distinct immediate responses to stepwise EF depletion, we grouped DEGs into five gene clusters based on their response to increasing dTAG^V^-1 ligand concentrations (Fig. 3E, Supplementary Table S5). Gene clusters G1 and G2 comprise EF-repressed targets that were upregulated upon dTAG^V^-1 treatment. G1 genes exhibited a gradual increase in expression from the lowest to the highest dTAG^V^-1 concentration. In contrast, genes in cluster G2 exhibited a steep upregulation in response to dTAG^V^-1 concentrations up to 15 nM, reaching a plateau at higher concentrations. These two EF-repressed clusters showed a strong enrichment in genes annotating to TNFα signaling, hypoxia and EMT (Supplementary Fig. S4E). For EF-activated targets, we observed three patterns: genes in the smallest cluster G3 exhibited a steep decline in expression starting at the lowest dTAG^V^-1 concentration (1.5 nM; ∼14% EF reduction) continuing up to 15 nM (∼57% EF reduction) (Fig. 3E). Interestingly, at higher dTAG^V^-1 concentrations, where EF levels approached near-complete depletion, a paradoxical upregulation in gene expression was observed. In contrast, genes in G4 and G5 decreased in expression upon gradual EF depletion. While the expression of G4 genes remained unchanged up to a dTAG^V^-1 concentration of 5 nM (∼35% EF reduction), G5 genes displayed a decline in expression already at 1.5 nM (∼14% EF reduction). Notably, cluster G5 includes several well-characterized GGAA microsatellite-driven genes such as *BCL11B*, *CCND1*, *NKX2-2*, *NPY1R, NR0B1* and *POU3F1* (6,44). In summary, we have identified a set of 810 genes with distinct sensitivities to acute EF dosage change, elucidating distinct EF dosage dependent regulatory mechanisms and pathways.

### Distinct architecture of EF chromatin binding regions determines pattern of response to EF modulation

The differences in the acute responses of individual gene expression clusters to gradual EF depletion may result from variable EF binding affinities to genomic regions with distinct chromatin architectures and from functional interactions with other transcription co-factors. We therefore conducted genome-wide profiling of EF-bound chromatin regions and associated histone landscapes in parental A673 cells and EF-dTAG clone A2.2 using CUT&RUN with antibodies directed to the FLI1 C-terminus, H3k27ac, H3k27me3, and H3k4me3. We identified a total of 12,370 consensus EF binding peaks, supported by replicates in either parental A673 or A2.2 cells (Supplementary Fig. S5A, Supplementary Table S1). The majority of EF binding was observed in promoters (44.5%) characterized by H3K27ac and H3K4me3 chromatin marks, followed by intronic (28.5%) and distal intergenic (20.4%) regions marked by H3K27ac enhancer marks (Supplementary Fig. S5B-C). In line with prior findings, FLI1-binding peaks showed significant depletion in regions marked by the repressive H3K27me3 mark (Supplementary Fig. S5C) (12).

We scanned the EF-bound chromatin regions for occurrences of transcription factor binding motifs from the JASPAR2020 core collection (45) and identified eight peak clusters based on recurrent motif associations (Fig. 4A-C, Supplementary Table S6). For example, the largest cluster, P1, contained 4160 peaks that were predominantly localized to promoter regions and enriched with binding motifs for Krüppel-like and other zinc finger transcription factors (KLF15/16, SP9, MAZ, and ZNF148). Peak cluster P3 (n=2079) showed significant enrichment for canonical monomeric ETS binding motifs. A distinct cluster, P5 (958 peaks), was uniquely characterized by a high enrichment of EF binding GGAA repeat motif matches. These peaks were wider and had a higher signal than those in other clusters and were primarily localized to intergenic regions. In addition, they were most sensitive to low dTAG^V^-1 concentrations as measured by fold change (Supplementary Fig. S5D).

**Figure 4.**
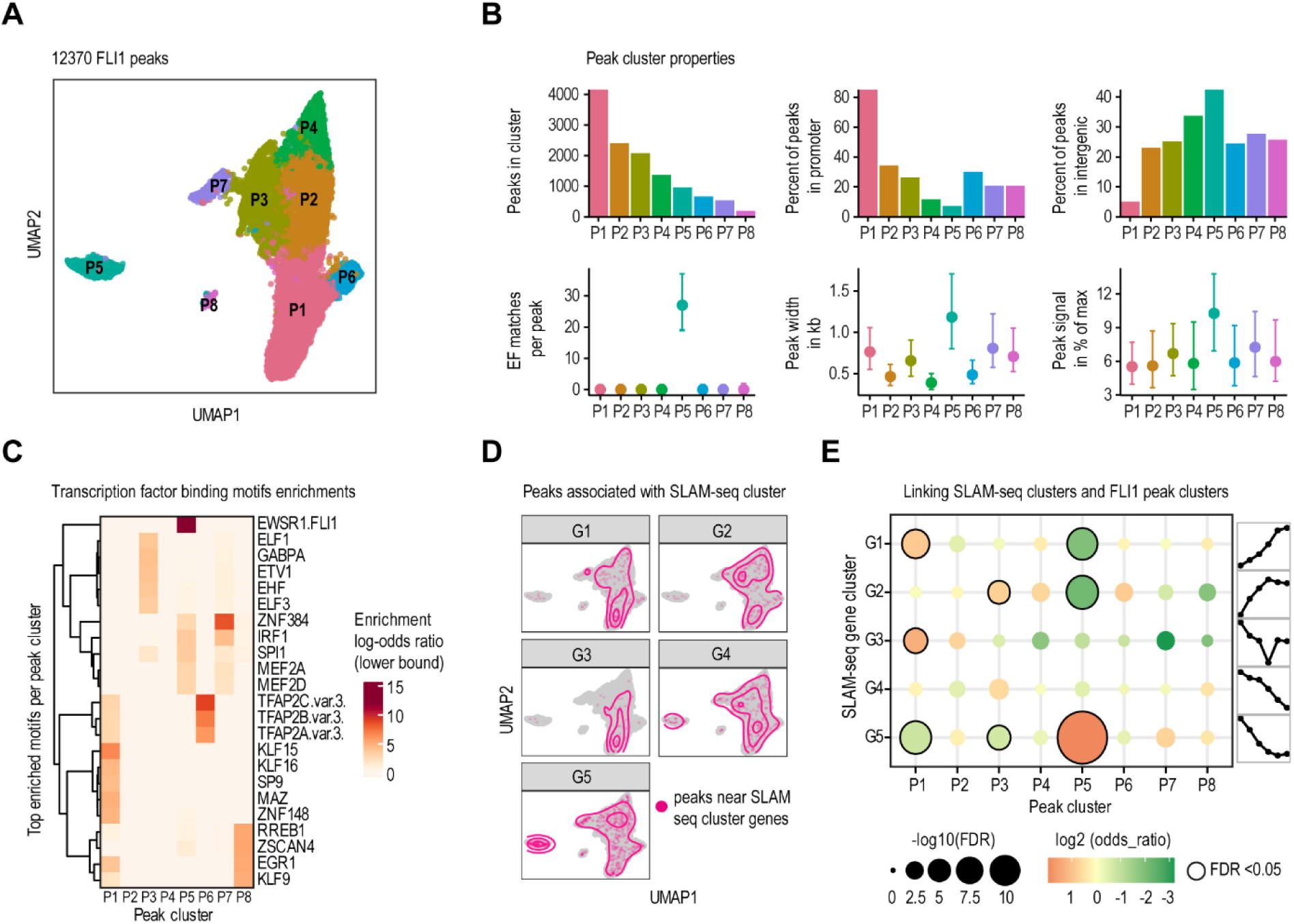
Profile of EF-bound chromatin regions and its association with immediate response genes. **A**, UMAP plots summarizing the results of a genome-wide analysis of EF-bound chromatin regions with CUT&RUN. Each point indicates one peak. A total of 12,370 consensus EF binding peaks were identified, supported by replicates in either parental A673 or A2.2 cells. JASPAR2020 ^45^ human core transcription factor binding motif matches were counted for every peak and the Manhattan metric was used to define peak-to-peak distances. Clustering was done with the Louvain method ^75^ and eight distinct EF-binding peak clusters were identified. **B**, For each peak cluster, the total number of peaks, their overlap with promoters or intergenic regions, number of matches to the EF binding motif (matrix profile MA0149.1), width, and CUT&RUN signal intensity (MACS2 reported peak signal rescaled to [0, 100] per sample, replicates averaged were summarized (median and interquartile range shown as points and ranges). **C**, Transcription factor binding motif match enrichments were calculated for every peak cluster, displaying the top five motifs. Motifs were filtered (one-sided Fisher’s exact test; FDR < 0.01, lower bound of 95% confidence interval of log_2_ odds ratio > 2) and ranked by the log_2_ odds ratio. **D**, Density contour plots overlaid on the UMAP plot from panel (**A**), indicating association of peaks with genes responsive to EF degradation. Every peak was linked to its closest response gene (out of the 810 differentially expressed SLAM-seq genes, cp. Fig. 3E) within 150kb distance. **E**, Cluster-specific enrichments were calculated for each EF-dependent gene cluster (G1-G5) and each EF-binding peak cluster (P1-P8) using two-sided Fisher’s exact test. Circle size and color indicate the FDR and odds ratio, respectively. Significant associations (FDR < 0.05) are highlighted with a solid border.

To link EF binding regions with immediate response genes from our SLAM-seq analysis, we mapped the EF-binding peaks to the closest transcription start sites (≤150 kb upstream or downstream) of actively expressed genes and tested for association between gene clusters and peak clusters (Fig. 4D-E). We found that EF-repressed genes in clusters G1, G2 (= increased expression with higher dTAG^V^-1) were significantly depleted near P5 peaks, i.e., peaks with intergenic GGAA microsatellites (Fig. 4E). Instead, genes within cluster G1 were associated with the promoter- and KLF-linked peak cluster P1. Interestingly, EF-repressed genes with the highest sensitivity to EF depletion (G2) were enriched near ETS-linked peak cluster P3. In contrast, EF-activated genes in G5 were significantly enriched near GGAA-microsatellite-linked peaks in cluster P5, and depleted near peak clusters P1 and P3, suggesting a distinct gene regulatory mechanism compared to EF-repressed genes.

### Sustained activation of EMT-associated transcriptional programs upon transient EF modulation

After establishing the acute response of EwS cells to gradual EF depletion, we investigated the molecular and phenotypic consequences of prolonged EF dosage modulation. A2.2 cells were treated with the dTAG^V^-1 gradient for 7, 14, 21, and 28 days, and B3.1 cells were treated for 21 days. To mimic dynamic EF fluctuations, a subset of cells underwent a 7-day washout of the ligand following the treatment period. For each condition, samples were collected for RNA-sequencing. Flow cytometry analysis demonstrated that the EF gradient remained stable across 7, 14, 21, and 28 days of dTAG^V^-1 treatment in EF-dTAG A2.2 clone (Supplementary Fig. S6A). Upon washout, baseline EF expression levels were restored within 16 hours (Fig. 1D) and remained stable at the end of the 7-day washout period (Supplementary Fig. S6B). Principal component analysis of the RNA-seq data demonstrated a clear separation between dTAG^V^-1-treated samples and untreated controls (Fig. 5A and Supplementary Fig. S6C), with gene expression changes correlating with both dTAG^V^-1 concentration and treatment duration (Fig. 5B and Supplementary Table S7). Enrichment analysis revealed EMT to be the top-significant pathway irrespective of the dTAG^V^-1 concentration or treatment durations in both the EF-dTAG clones (Fig. 5C and Supplementary Fig. S6D). Washout samples were clearly separated from treated samples, but the complete recovery of EF levels only partially reversed the observed gene expression changes. The largest number of dysregulated DEGs were seen after prolonged exposure to higher dTAG^V^-1 concentrations (Fig. 5D, Supplementary Fig. S6E left and Supplementary Table S7). Across all treatment durations and concentrations, most of the sustained effects could be attributed to EF-repressed genes, which remained upregulated after rescue of fusion protein expression (Fig. 5D and Supplementary Fig. S6E left). EMT, TNF-alpha signaling, hypoxia, and myogenesis-related gene sets were significantly enriched, displaying a consistent response pattern that remained largely independent of dTAG^V^-1 concentration in cells treated for 21 or 28 days, followed by a 7-day EF rescue in both EF-dTAG clones (Fig. 5E and Supplementary Fig. S6E right). A total of 106 genes were consistently upregulated in both EF-dTAG clones following EF rescue after 21 days treatment at any dTAG^V^-1 concentrations. Notably, 23 of these genes were associated with the EMT gene set from the MSigDB Hallmark 2020 pathways (Fig. 5F). These results demonstrate that EF modulation for at least 7 days persistently alter the EwS transcriptome, fostering an EMT transcriptional program.

**Figure 5.**
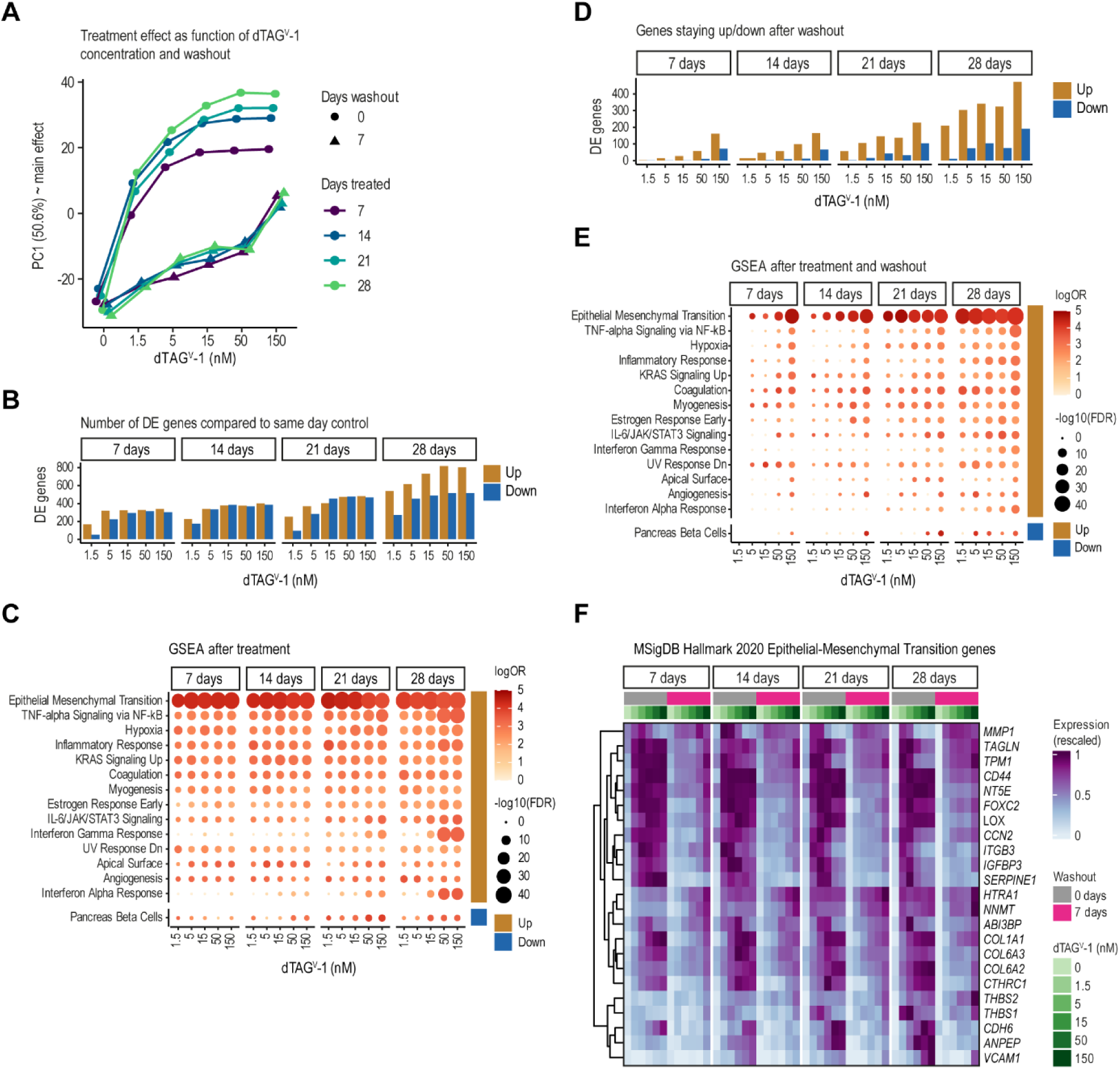
Transcriptional effects of prolonged and transient EF modulation. **A**, Principal component analysis (PCA) of 1,455 highly variable genes (gene expression with standard deviation > 0.5 after normalization with DESeq2^74^ vst function and averaging two replicates), highlighting the primary transcriptional effects of prolonged and transient EF dosage modulation (RNA-seq). The plot shows the value of principal component 1 (PC1), the main treatment effect, at different dTAG^V^-1 concentrations. Point shape distinguishes conditions without (circle) and with 7-day washout (triangle), and color indicates the number of days of dTAG^V^-1 treatment. **B**, Number of DEGs following dTAG^V^-1 gradient treatment compared to untreated controls at the same day (DESeq2; FDR < 0.05, absolute log_2_ fold change > 2). **C**, Functional enrichment analysis of DEGs after treatment, showing MSigDB Hallmark 2020 pathways that are significantly enriched in any of the dTAG^V^-1 treatments (hypergeometric test; FDR < 0.001, log_2_ odds ratio > 3; background: all DESeq2 tested genes). Circle size and color indicate the FDR-adjusted p-value and the odds ratio, respectively. **D**, Number of DEGs following treatment and subsequent washout compared to untreated control at the same day (DESeq2; FDR < 0.05, absolute log_2_ fold change > 2). **E**, Functional enrichment of DEGs after dTAG^V^-1 treatment followed by washout (cf. **C**). **F**, Heatmaps depicting genes from the MSigDB Hallmark 2020 Epithelial-Mesenchymal Transition pathway that are upregulated after a three-week treatment and persist following ligand washout for at least one dTAG^V^-1 concentration in clone A2.2 and B3.1. Expression values are averages of replicates after normalization (cf. **A**). Conditions on the x-axis are ordered by treatment duration, washout status, dTAG^V^-1 concentration, as indicated above the heatmaps.

### EwS cells retain increased metastatic potential upon transient EF modulation

To validate the transcriptomic findings at a functional level, both EF-dTAG clones were treated with gradient concentrations of dTAG^V^-1 over 7 or 21 days, followed by a 7-day ligand withdrawal. Colony formation, cell migration, and invasion assays were then performed, as previously described (Fig. 2). The washout of dTAG^V^-1, after a 7-day pre-treatment with the ligand, fully rescued the EF gradient-dependent migration phenotype, restoring it to the level observed in untreated cells (Supplementary Fig. S7A). However, following a 21-day pretreatment with dTAG^V^-1 ligand, withdrawal for 7 days was insufficient to fully reverse the migration phenotype. This incomplete recovery was observed at dTAG^V^-1 concentrations of 5–150 nM for clone A2.2 and 5– 15 nM for clone B3.1 (Fig. 6A, Supplementary Fig. S7B). Prolonging the ligand washout period to two weeks also failed to fully revert the migratory potential of 21 days dTAG^V^-1 treated EF-dTAG clones to that of untreated controls (Supplementary Fig. S7C-D). Likewise, organ-on-a-chip invasion assays demonstrated a significant retention of invasive properties of 21 days dTAG^V^-1 treated EF-dTAG clones after 7 days of EF recovery in the organoplate (Fig. 6B and Supplementary Fig. S7E). Consistent with the consequences of short term dTAG^V^-1 treatment, soft-agar colony formation of EF-dTAG clones decreased with gradual EF modulation for 21 days and was completely abolished at dTAG^V^-1 concentrations above 5 nM (Fig. 6C and Supplementary Fig. S7F). However, when these 3-week pre-treated cells were seeded and subsequently maintained in absence of dTAG^V^-1 throughout the soft-agar period, a significant recovery of colony forming ability was observed (Fig. 6C and Supplementary Fig. S7F). These findings indicate that transient EF modulation can condition cells to maintain migratory and invasive phenotypes, while preserving anchorage-independent growth.

**Figure 6.**
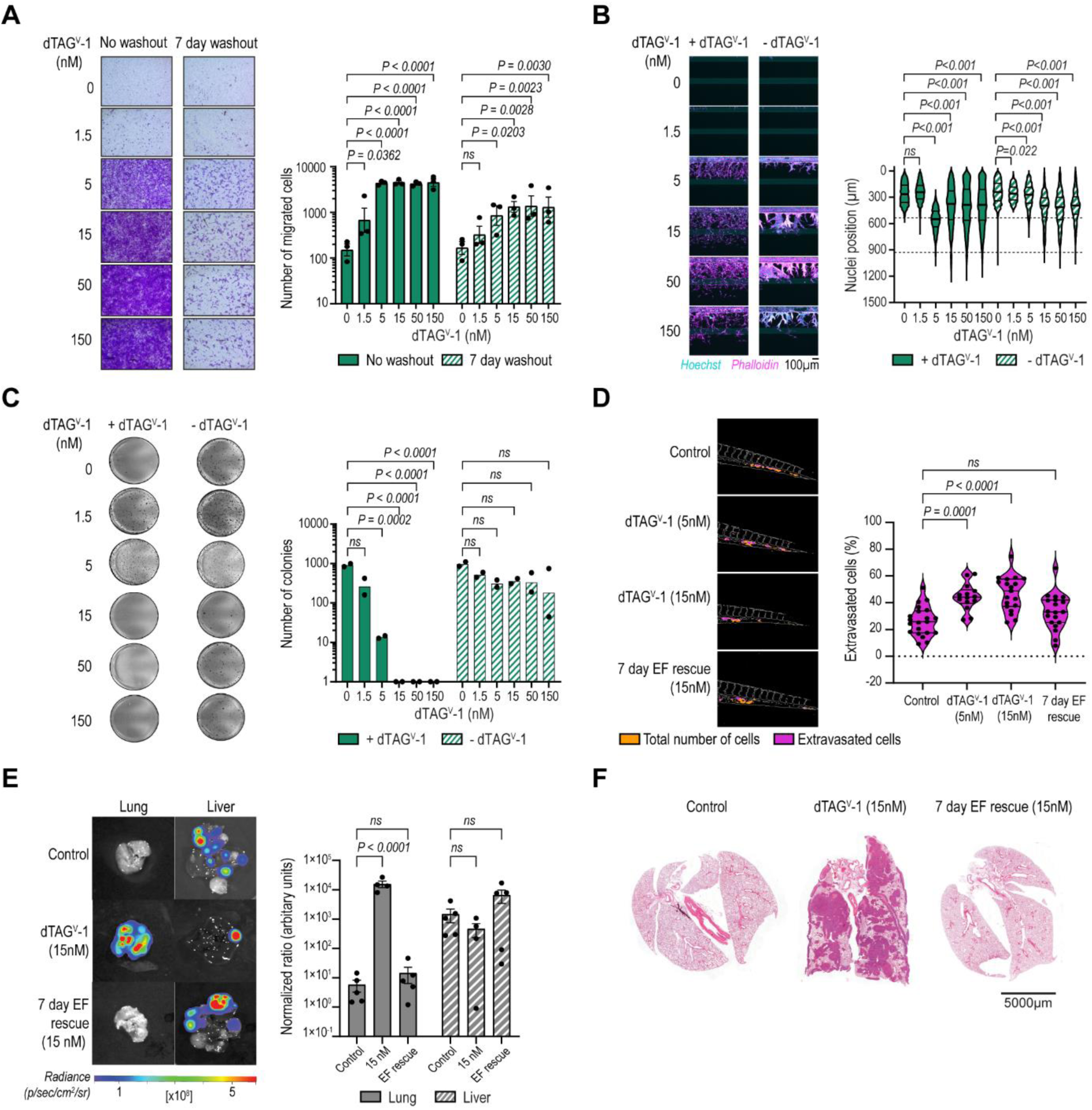
In vitro and in vivo assessment of metastatic effects following transient EF modulation. **A**, Transwell migration assays showing migrated A2.2 cells pre-treated with dTAG^V^-1 for 21 days at the indicated concentrations or pre-treated for 21 days followed by a 7-day washout. Representative images from one of three independent experiments are shown (left). Migrated cell counts are presented as mean ± SEM after log10 transformation (right; *n* = 3). **B**, Organoplate invasion assays depicting cell invasion into collagen I ECM 7 days after seeding A2.2 cells treated with dTAG^V^-1 for 21 days at the indicated concentrations, with or without further dTAG^V^-1 treatment. Representative images from one of two independent experiments are shown (left). Nuclei positions along the Y-axis are shown, with phase guides indicated by dotted lines at 535 µm and 930 µm (right; *n* = 2). **C**, Soft agar assays showing colonies formed by A2.2 cells three weeks after seeding 21 days dTAG^V^-1 treated cells at indicated concentrations with or without further dTAG^V^-1 treatment. Representative images from one of two independent experiments are shown (left). Colony counts are presented as mean (*n* = 2) following log10+1 transformation. **D**, Zebrafish extravasation assays 3 days post injection of cells treated with dTAG^V^-1 for 21 days at the indicated concentrations, or 21 days of 15 nM dTAG^V^-1 followed by a 7-day washout. Representative images from one of the technical replicates are shown (left). Extravasation data are presented as a percentage of extravasated cells (right, *n* = 21 for control, *n* = 14 for 5 nM, *n* = 19 for 15 nM and EF rescue). **E**, Endpoint In-vivo Imaging System (IVIS) imaging of lungs and livers from mice tail vein injected with A2.2 cells treated with 15 nM dTAG^V^-1 for 21 days, with or without a 7-day washout, with no further dTAG^V^-1 treatment. Representative images from one of the technical replicates are shown (left). Quantification of organ-wide luciferase intensity from tumors in the lungs and livers, normalized to whole-body luminescence of their respective conditions at day 0 (right, *n* = 5 mice/group, *n* = 4 mice for 15 nM). **F**, H&E-stained lung sections from the conditions shown in panel e. Representative images from one of the technical replicates are shown. *P*-values for panels **A**, **B**, **C**, and **E** were calculated using two-way ANOVA with Dunnett’s post-hoc multiple comparisons test. *P*-values for panel **D** were calculated using One-way ANOVA with Dunnett’s post-hoc multiple comparisons test. *ns*, not significant.

We hypothesized that these phenotypic changes in cells were linked to an increased capacity to metastasize *in vivo*. To investigate this hypothesis, we first examined the extravasation properties of A2.2 cells following partial EF depletion and subsequent rescue using a zebrafish model. A2.2 cells were treated with 5 nM or 15 nM dTAG^V^-1 for 3 weeks before being intravascularly injected into zebrafish larvae with no further dTAG^V^-1 ligand provided, thus allowing recovery of the fusion protein. Additionally, we included an EF rescue condition by culturing the 15 nM dTAG^V^-1 pre-treated cells without ligand for 7 days prior to injection. At 3 days post injection, a significant increase in the number of extravasated tumor cells was observed in both the 5 nM and 15 nM dTAG^V^-1 pre-treated groups compared to untreated controls (Fig. 6D). In contrast, the 7-day EF rescue condition did not exhibit significant differences in extravasation levels compared to untreated controls.

Next, we injected luciferase-expressing A2.2 cells with or without 21-day pre-treatment (15 nM dTAG^V^-1) into the tail vein of NSG mice. Again, we included an EF rescue condition where dTAG^V^-1 treatment was ceased 7 days prior to injection (Supplementary Fig. S7G). Animals were subsequently monitored for 32 days in the absence of dTAG^V^-1 to allow *in vivo* recovery of the EF fusion protein. Independent of treatment, all mice injected with EF-dTAG cells showed a comparable tumor burden in the liver, but cells pre-treated with 15nM dTAG^V^-1 exhibited a significantly increased lung metastatic burden compared to untreated controls (Fig 6E). These findings align with observations from the zebrafish extravasation model and suggest that transient EF protein modulation promotes tumor cell seeding and overt colonization of lung tissue in mammals. Histological analysis of H&E-stained lung sections confirmed enhanced metastatic burden in mice injected with dTAG^V^-1 pre-treated cells (Fig. 6F). Interestingly, despite retaining an elevated EMT transcriptional signature and increased *in vitro* migratory and invasive potential, the 7-day EF recovery condition did not significantly result in increased lung metastasis following tail vein injection (Fig 6E). Together, these results suggest that after transient EF modulation, recovery of the fusion protein restores the extravasation properties of EwS cells and their ability to develop clinically relevant lung metastases, while their invasive and migratory propensity remains increased.

### Signature genes of transient EF modulation are associated with adverse prognosis in EwS patients

We demonstrated that a transient reduction in EF levels persistently activates an EMT transcriptional signature in EwS cells associated with increased migratory and invasive behavior in experimental models. To evaluate the clinical relevance of this gene expression signature, we analyzed a cohort of 196 clinically well-annotated EwS patients (46–49). Microarray-based gene expression data were available for 85 persistently upregulated genes in both EF-dTAG clones following EF rescue after 21-day treatment at any dTAG^V^-1 concentrations. Notably, high expression of 23 of these genes were significantly associated with poor overall survival (Fig. 7A and Supplementary Table S8). Examples for prognostic genes include *LOX, NAV2, PDGFRA, THBS2, CDCP1, SEMA3C,* which have been shown to promote metastasis in various cancers (50–55) (Fig. 7B). Together, these data suggest that prolonged, but transient modulation of EF leads to persistent dysregulation of a subset of genes associated with an aggressive disease phenotype.

**Figure 7.**
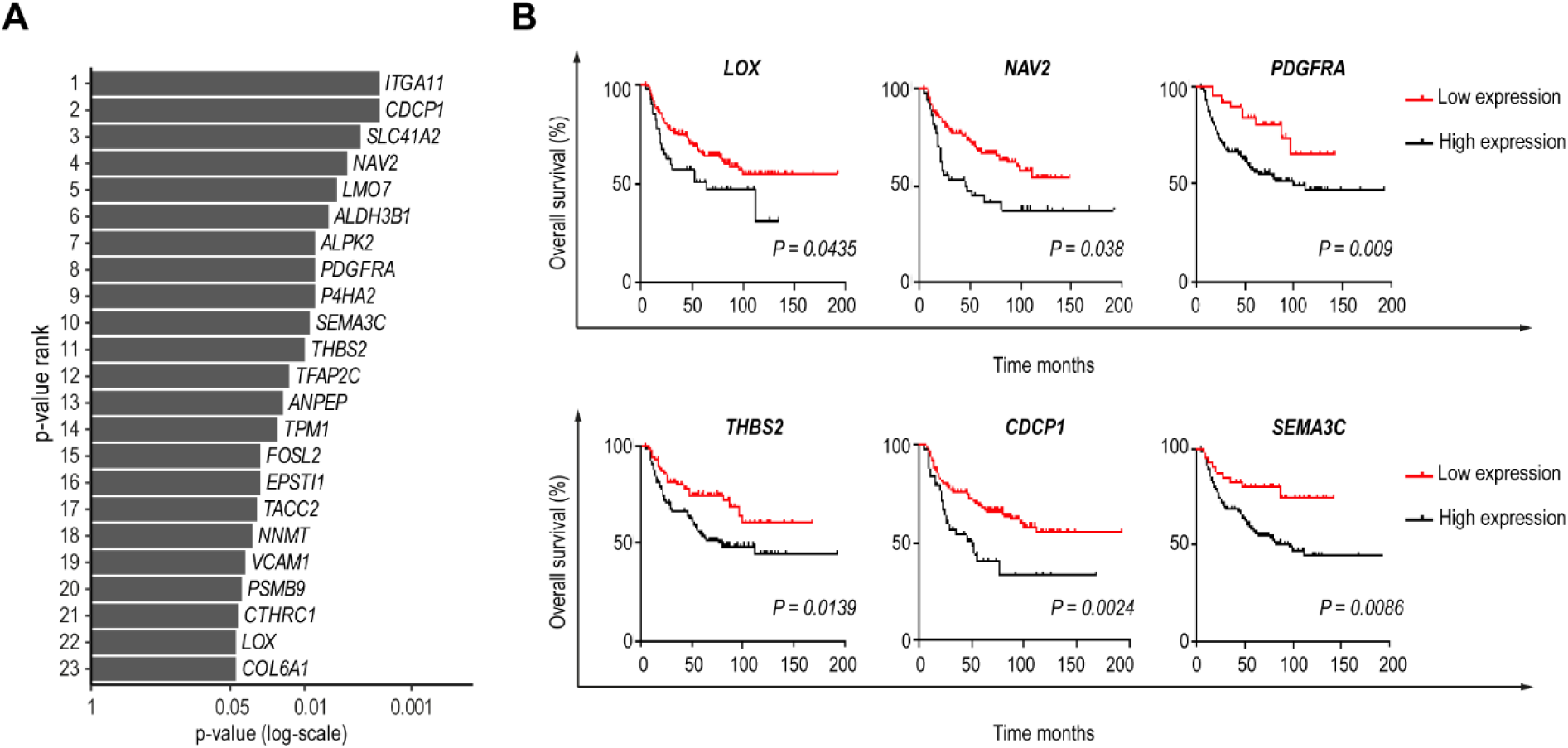
Survival analysis of dysregulated genes resulting from transient EF modulation. **A**, Rank plot depicting a subset of genes persistently upregulated in EF-dTAG clones after 21 days of dTAG^V^-1 treatment followed by a 7-day washout. The genes shown are those with a significant association (Log-rank (Mantel-Cox) test, p-value < 0.05) with poor overall survival when highly expressed in a cohort of EwS patients (*n* = 196). **B**, Kaplan-Meier survival curves for selected representative metastasis related genes identified in panel **A**, demonstrating their prognostic relevance.

## DISCUSSION

In this study, we investigated the direct relationship between EF oncoprotein dosage and EwS cellular phenotype at both molecular and functional levels. Knock-in of a fluorescent and a degron tag at the FLI1 carboxy-terminus was well tolerated in the primary non-metastatic tumor-derived A673 cell line with minimal transcriptomic variation observed when compared to single cell clones from unedited parental A673 cells. Although the EF carboxy-terminus was previously shown to play a role in protein interactions involved in cooperative DNA binding with other transcription factors (56), we did not observe major differences in genome-wide EF chromatin occupancy between the tagged and the untagged oncoprotein in A673 cells. However, our failed attempts to establish a similar model in other EwS cell lines suggest that tolerance to the carboxy-terminal EF modification may be context-dependent.

A key advantage of our degron model is its stability and reproducibility in revealing distinct phenotypic states associated with precisely defined and sustained EF levels. Using this model, we demonstrate for the first time that already a minor reduction in EF levels by as little as 14% was sufficient to induce a migratory and invasive phenotype in EF-dTAG cells which peaked at an approximately 57% EF reduction. However, upon further EF depletion (84-93%) migratory and invasive capabilities markedly decreased, suggesting that a critical threshold of EF is required for sustaining pro-metastatic potential. Remarkably, even minimal EF modulation (around 14% EF reduction) decreased soft-agar colony-forming ability, while no effects on cell proliferation were observed. Our study identifies precise EF thresholds that govern EMT-like transitions, elucidating insights previously suggested by crude EF knockdown experiments (19,23,24). Furthermore, we extend and refine the “Goldilocks principle” of EF dosage-dependent regulation initially proposed for EwS cell growth (57) to include pro-metastatic tumor cell properties.

To understand the molecular basis of EF level-dependent phenotypic plasticity, we monitored the acute and direct transcriptional response to graded EF depletion hypothesizing that EF affinity to distinct target gene sets may vary in a concentration-dependent manner. Indeed, we found clusters of genes with distinct response patterns to graded EF depletion. We found GGAA microsatellite-driven EF-activated genes (G5) to be the most sensitive to even subtle (∼14%) EF modulation, with almost complete loss of expression at an EF protein reduction of 57%. While ETS transcription factor binding sites usually comprise a specific sequence of 6 to 8 base-pairs with a central GGA/T motif, EF has been shown to bind to arrays of simple GGAA repeats as multimers by a mechanism that involves biomolecular condensation via the EWS amino terminal low-complexity domain (58,59). Endogenous EF concentrations in EwS cell nuclei have been estimated to be in the range of 200 nM (58). *In vitro*, lowering the EF protein concentration from 250 to 100 nM resulted in a marked drop in GGAA repeat-associated condensate formation (59), which is in line with our *ex vivo* observation that already minor decreases in endogenous EF thresholds lead to a loss of GGAA microsatellite-associated target gene activity. In sharp contrast, EF-activated gene expression cluster G4 was much more robust and only poorly sensitive to acute EF loss of up to 35%, followed by an EF concentration-dependent gradual decrease of gene expression, with complete loss observed only upon after near-complete EF depletion. Lack of enrichment for specific transcription factor motifs in EF-binding peaks associated with this group of genes may suggest involvement of other unidentified EF co-factors or an early indirect mechanism of gene regulation.

Among the EF-repressed genes, those characterized by the presence of monomeric canonical ETS binding sites near EF-bound chromatin (G2) showed the highest sensitivity to subtle EF modulation, with maximum de-repression occurring at ∼57% EF reduction. Of note, these genes completely lacked association with EF-bound GGAA microsatellites. This pattern is consistent with previous reports suggesting two distinct binding site preferences of EF in EwS, where EF-activated sites typically contain four or more consecutive GGAA repeats, whereas EF-repressed sites are often linked to non-repetitive, canonical ETS motifs (5,7,60). However, unlike studies that mainly associate gene activation with EF binding at GGAA microsatellites within promoter regions (6,60), our data indicate a predominant association with intergenic regions consistent with findings that GGAA microsatellites act as potent distal enhancers in EwS (7,61).

The lack of enrichment for canonical ETS and E2F binding motifs among EF-activated DEGs in SLAM-seq experiments came as a surprise, as previous studies on EF chromatin binding have shown cooperative binding of EF with E2F3 in the promoters of E2F-regulated cell cycle genes (4). Knockdown of EF has been shown to result in the replacement of activating E2F3 by the repressive E2F4 family member (62). Although we observed reduced EF binding to canonical ETS binding motifs already at 3 hours of dTAG^V^-1 treatment (EF binding peak cluster P3, Fig. 4E), we speculate that E2F3 replacement may require more time to affect the expression of cell cycle genes. Indeed, after 24 hours of EF modulation, E2F target and G2/M checkpoint genes were most significantly enriched at EF reductions of at least 57%, suggesting robustness of EF-driven E2F target gene regulation to subtle short-term variations of EF expression.

EF-repressed immediate response genes lacking both canonical ETS and GGAA microsatellite motifs represented the largest gene expression cluster (G1) and showed a graded response to stepwise EF depletion. A significant enrichment of binding motifs for Krüppel-like three-zinc finger transcription factors was found predominantly in promoters of G1 cluster genes. Among them was the transcriptional repressor KLF15, which has previously been implicated in the core regulatory circuitry of EF-interacting transcription factors, activated through EF binding to a GGAA microsatellite super-enhancer (63) and expressed as part of gene expression cluster G5. Its potential role in EF-mediated transcriptional repression remains to be established. Additionally, binding motifs for zinc finger transcription factors MAZ and ZNF148 were found enriched in promoter-associated EF binding peaks of G1 cluster genes. So far, nothing is known about functional interactions between EF and any of these candidate EF-interacting transcription factors. Notably, G1 cluster genes were strongly enriched for EMT-related pathways, suggesting a potential involvement of these co-factors in driving the migratory and invasive phenotypes observed with gradual EF depletion.

A particularly intriguing group of acute EF response genes, which also include *MYC* and its target genes (i.e., *TFB2M*, *HNRNPC* and *SNRPD1*), exhibits a bimodal expression pattern that is dependent on EF thresholds (G3). These genes display a sharp decline in expression following subtle EF reductions, similar to the behavior observed in the GGAA microsatellite-driven cluster G5, with complete loss of expression occurring at residual EF levels of about 43%. However, upon further EF depletion the expression of these genes resurges. A potential explanation for this increase of gene expression at very low EF concentrations may be re-activation by an EF-suppressed transcription factor which remains to be defined. EF binding peaks in the vicinity of these genes were enriched in the same transcription factor binding motifs (P1) as G1 cluster genes and did not show a particular preference for either promoter or intergenic regions. Of note, bimodal expression response of G3 cluster genes to EF depletion inversely correlated with variations in migratory and invasive phenotypes of EF-dTAG cells. These phenotypes peaked at an EF threshold of 43% and declined as G3 cluster gene expression increased at higher dTAG^V^-1 concentrations. Although gene set enrichment analysis did not reveal specific functional annotations for this cluster, the G3 genes may represent critical regulators of metastasis, warranting further investigation in future studies.

Our findings indicate that EwS tumor phenotype is not solely dependent on the binary presence or absence of EF but is instead influenced by EF dosage levels. The downstream effects of direct EF inhibition, as well as the mechanisms of resistance that may arise, remain incompletely understood. To address these gaps, we leveraged the capability of the dTAG degron system to rapidly re-establish EF expression after dTAG^V^-1 ligand withdrawal, following transient dTAG^V^-1-mediated graded EF degradation. This system allowed us to simulate an EF-directed therapeutic approach and examine the dynamics of EF target gene recovery and its impact on tumorigenesis. Notably, transient EF degradation for 1-4 weeks, followed by EF re-expression for 1 week, led to a sustained transcriptional dysregulation of mostly EF-repressed EMT-related genes, irrespective of EF levels. This result implicates epigenetic fixation of EF target gene dysregulation although the exact mechanism behind this observation remains to be defined. Strikingly, approximately 27% of genes remaining upregulated post-EF recovery were associated with poor patient prognosis. Consistent with these findings, *in vitro* assays revealed a significant retention of an elevated migratory and invasive phenotype and partial retention of colony forming ability in soft-agar following 7-day EF rescue after three weeks of EF modulation. However, while injection of dTAG^V^-1 pre-treated EwS cells led to increased extravasation in zebrafish and lung metastasis in a mouse tail vein injection model upon EF restoration during the *in vivo* incubation period, xenografting of already 7-day EF-rescued cells did not increase metastatic burden. These results suggest that extravasation is a rate-limiting factor in EwS metastasis, and may critically require EF modulation, corroborating with previous observations from a crude EF-knockdown model (19). Historically, transcription factors have been considered ‘undruggable’. However, recent advancements in small-molecule therapeutics have shown promise in preclinical and clinical cancer settings (64,65). For EwS, several agents have been identified that interfere with EF target gene expression, including cytarabine (66), YK-4-279 and TK216 (67,68), trabectedin (69), mithramycin and other mitralogues (70,71), midostaurin (72), low-dose actinomycin (73), shikonin (74), and HCI2509 (10). Most of these agents do not directly target EF protein expression but instead inhibit GGAA microsatellite-dependent transcriptional activation, with limited focus on their effects on the EF repressive signature. Our findings underscore the importance of this repressive signature, which plays a critical role in EMT and metastasis, and demonstrate that it can be irreversibly disrupted by transient EF fluctuations. These observations have significant implications for single-driver oncoprotein-targeted therapies in EwS, where incomplete EF inhibition may unintentionally promote metastatic progression and contribute to relapse.

## METHODS

### Cell culture

The A673 EwS cell line derived from a primary tumor localized in muscle, was purchased from Cell Lines Service (CLS GmbH, Cat# 300454, RRID: CVCL_0080) and authenticated by short tandem repeat (STR) profiling at Microsynth. All cells were cultured in complete DMEM medium (Gibco, Cat# 31966-021) supplemented with 10% fetal calf serum (FCS, Gibco, Cat# A5256701) and 1% penicillin-streptomycin (PAN Biotech, Cat# P06-07100) at 37°C and 5% CO_2_. Cells were passaged at a 1:10 ratio every 3-4 days, or upon reaching confluency, using Accutase (PAN Biotech, Cat# P10-21100). Regular mycoplasma testing was performed using the MycoAlert Mycoplasma detection kit (Lonza, Cat# LT07-318).

### Plasmids and cloning materials used in this study

All plasmids and primers used in this study are summarized in Supplementary Table S9. To facilitate the cloning of the repair template plasmid for endogenous knock-in of A673 cells, primers were designed to amplify 500 bp homology arms corresponding to genomic sequences located immediately 5′ and 3′ of the sgRNA cleavage sites. Another set of primers was used to PCR amplify the mNeonGreen-HA-FKBP12^F36V^ fragment from the pCRIS-PITChv2-internal-mNG-dTAG plasmid (provided by Georg Winter’s lab(75)). All amplified fragments were verified for single nucleotide polymorphisms (SNPs) by Sanger sequencing before being assembled into an EcoRV-digested pUC57 plasmid backbone (Genscript, Cat# SD1176) using the Gibson Assembly® Cloning Kit (NEB, Cat# E5510S). For *in vivo* mouse experiments, Luc2 expressing dual marker selection vector (pRP[Exp]-EGFP/Hygro-CAG>Luc2, VectorBuilder Cat# VB900120-5146vmm), obtained as a gift from Oscar Martinez Tirado’s lab (IDIBELL Biomedical Research Institute of Bellvitge), was transfected directly into parental A673 and A2.2 cells using the TransIT-X2® Dynamic Delivery System (Mirus Bio, Cat# MIR 6004) as per the manufacture’s protocol.

### CRISPR–Cas9 genome editing of A673 cells

A CRISPR-mediated knock-in approach was employed to tag endogenous EF in A673 cells with mNG fused to HA and dTAG. An sgRNA targeting the exon-intron junction of the last exon of the FLI1 C-terminal locus was designed using the CHOPCHOP tool (RRID:SCR_015723) (76) and selected based on proximity to the FLI1 stop codon and the highest MIT specificity score. The sgRNA sequence was ordered as an Alt-R CRISPR-Cas9 sgRNA (IDT Cat# 11-01-03-01, Supplementary Table S9). The cloned pUC57-based repair template plasmid was used to generate a single-stranded donor template with the Guide-it long ssDNA Production System v2 (Takara, Cat# 632666) using the primers listed in Supplementary Table S9 (Takara, Cat# 632666). For nucleofection, 1 million low-passage A673 cells were resuspended in nucleofector solution (Cell Line NucleofectorTM Kit V, Cat# VCA-1003) together with 2 µg of either sense or anti-sense ssDNA and 2 µL of 10.8 µM Alt-R Cas9 Electroporation Enhancer (IDT, Cat# 1075916). Concurrently, an RNP complex was formed by combining Cas9 protein (Alt-R S.p. HiFi Cas9 Nuclease V3, Cat# IDT1081060) with sgRNA at a 1:1.2 molar ratio in a total volume of 10 µL with DPBS (Gibco, Cat# 14190169). This mixture was incubated for 15 minutes at room temperature, then added to the cells for nucleofection using the Amaxa Nucleofector 2b (program x-001). Cells were then transferred to a 6-well plate containing 1 mL DMEM medium without antibiotics, supplemented with 15 µL of Alt-R HDR Enhancer (3 mM stock solution, IDT, Cat# 1081072). After 24 hours, the medium was replaced with a complete growth medium for cell recovery. Single cells expressing mNeonGreen were sorted by fluorescence-activated cell sorting (FACS) using a BD FACSAria^TM^ Fusion (RRID:SCR_025715) and plated into 96-well plates.

### Validation of Knock-In

Knock-in validation was initially confirmed through two-step RT-PCR using cDNA derived from the modified cells to specifically assess integration at the transcript level. Total RNA was extracted with the RNeasy Mini Kit (Qiagen, Cat# 74106) according to the manufacturer’s protocol, including on-column DNase digestion to remove genomic DNA contamination. First-strand cDNA synthesis was performed using 1 µg of RNA, M-MLV Reverse Transcriptase (Promega, Cat# M1705), and Oligo(dT)15 primers (Promega, Cat# C1101). PCR amplification was performed using the resulting cDNA as a template, knock-in validation primers (Supplementary Table S9), and Q5® Hot Start High-Fidelity DNA Polymerase (New England Biolabs, Cat# M0493L) according to the manufacturer’s guidelines. PCR products were analyzed using 0.8% TAE gel electrophoresis alongside a 1 kb DNA ladder (Promega, Cat# G5711) to confirm successful integration of the tags into the EF locus. To further validate correct integration and expression of the mNG-HA-dTAG construct at the endogenous EF locus, we conducted Western blot analysis, confocal microscopy, and flow cytometry.

### dTAG^V^-1-mediated EF protein gradient

To establish an EF protein gradient, EF-dTAG clones were treated with increasing concentrations of dTAG^V^-1 ligand (Bio-Techne, Cat# 7374). Serial dilutions were prepared from a 10 mM stock to obtain micromolar working stocks, which were further diluted 1000-fold to achieve final dTAG^V^-1 concentrations of 1.5 nM, 5 nM, 15 nM, 50 nM and 150 nM. Cells were treated with these concentrations for different time intervals as indicated in the text. DMSO was used as vehicle control. For ligand washout, cells were washed twice with DPBS (Gibco, Cat# 14190-094), detached using Accutase, and transferred to new culture flasks to allow full recovery of EF expression.

### Flow cytometry

Flow cytometry was employed to evaluate EF expression upon dTAG^V^-1 treatment for each experiment, using mNG as a readout. Cells were harvested, washed, and resuspended in Clini MACS buffer (0.5 % BSA, 2 mM EDTA in DPBS). DAPI was added at a final concentration of 10 µg/mL (DAPI Staining Solution Miltenyi Biotec, Cat# 130-111-570) to exclude dead cells. Fluorescence intensity of mNG was measured using an BD LSRFortessa Cell Analyzer (RRID:SCR_018655) and data analysis was performed with FCS Express 7 (De Novo software, RRID:SCR_016431).

### Antibodies

The following primary antibodies were used for Western blot (WB) or CUT&RUN (C&R): FLI1 (Abcam Cat# ab133485, RRID:AB_2722650, C&R 1:30), IgG (Epicypher Cat# 13-0042, RRID:AB_2923178, C&R 1:50), H3K4me3 (Epicypher Cat# 13-0041, RRID:AB_3076423, C&R 1:50), H3K27ac (Cell Signaling Technology Cat# 8173, RRID:AB_10949503, C&R 1:50), H3K27me (Cell Signaling Technology Cat# 9733, RRID:AB_2616029, C&R 1:50), Tubulin (Millipore Cat# CP06, RRID:AB_2617116, WB 1:5,000).

### Western blot

Cells were directly lysed in 1× Laemmli buffer (2% sodium dodecyl sulfate, 10% glycerol, 5% 2-mercaptoethanol, 0.002% bromophenol blue, 0.0625 M Tris-HCl, pH 6.8) and boiled for 5 min at 95°C. Lysates were subjected to SDS-PAGE on 8% polyacrylamide gels, and proteins were transferred onto 0.45µm nitrocellulose membranes (Cytiva, Cat# 10600002) using a transfer buffer containing 20% methanol. Membranes were blocked with Intercept blocking buffer (LI-COR, Cat# 927-70001) for 1 hour at room temperature and then probed with primary antibodies overnight at 4°C. After three washes with TBS-T (200 mM Tris, 1500 mM NaCl, 0.1% Tween-20, pH 7.6), membranes were incubated with goat anti-rabbit DyLight 800 (Invitrogen, Cat# SA5-35571) or goat anti-mouse DyLight 680 (Invitrogen, Cat# 35518) IgG secondary antibodies diluted 1:10,000 for 1 hour at room temperature. Following three additional washes in TBS-T and rinsing with PBS, protein bands were visualized using the LI-COR Odyssey Classic Imager (RRID:SCR_023765) and LI-COR Image Studio Software version 5.2 (RRID:SCR_015795). Quantification of band intensities was performed using Fiji (RRID:SCR_002285) (77,78).

### Viability assays

Cell viability and caspase activity were assessed using the CellTiter-Glo Luminescent Cell Viability Assay (Promega, Cat# G7570) and Caspase-Glo 3/7 Assay (Promega, Cat# G8091) following the manufacturer’s protocols. One day prior to the experiment, cells were seeded in 96-well plates at a density of 10,000 cells per well. The following day, medium was replaced with corresponding gradient concentrations of dTAG^V^-1 ligand. After 48 hours of incubation at 37°C and 5% CO₂, an equal volume of 1:4 diluted CellTiter-Glo reagent in DPBS or Caspase-Glo 3/7 reagent was added to each well. Plates were incubated at room temperature for 10 minutes for CellTiter-Glo and 30 minutes for Caspase-Glo 3/7. Luminescence was measured using an EnSpire microplate reader (PerkinElmer). All experiments were performed in triplicate, with three technical replicates per condition. S4U cytotoxicity was measured for concentrations up to 2 mM over a time scale of 8 hours using Cell Titer Glo assay. S4U-containing medium was renewed after 4 hours. Even at the highest concentration, no decrease in cell viability was observed (Data not shown).

### Transwell migration assays

Cells were pre-treated with dTAG^V^-1 ligand for 18 hours followed by 4-hour starvation in serum-free DMEM medium in the presence of dTAG^V^-1 at indicated concentrations. Transwell inserts with a pore size of 8.0 µm (Corning, Cat# 3422) were placed into 24-wells containing 600μL complete medium. Subsequently, 200,000 cells were resuspended in 200μL DMEM medium containing 0.1% FCS and seeded into the upper chamber. Each condition was performed in duplicate. dTAG^V^-1 ligand was added to both the bottom and upper chamber, while DMSO served as the vehicle control. After 18-20 hours of incubation at 37°C, non-migratory cells on the upper surface of the membrane were removed using a cotton swab. Migratory cells that traversed the membrane were fixed with 4% formaldehyde solution (Sigma-Aldrich, Cat# F8775) and stained with 0.01% crystal violet in 0.25% methanol for 20 minutes. Images were taken at 10× magnification using Zeiss Axio Vert series Axiovert 200 inverted microscope (RRID:SCR_020915) equipped with an AxioCam and Zeiss Zen 2.3 Lite software (RRID:SCR_023747). Quantification was performed by counting cells in two fields of view per technical replicate using Fiji software.

### Organoplate invasion assays

Invasion assays were conducted using the Organoplate 3-lane 64 (Mimetas B.V.), which consists of 64 chips per plate, each containing 3 microfluidic channels (79). One day prior to cell seeding, Collagen I was prepared at a final concentration of 4 mg/ml (Cultrex 3-D Culture Matrix Rat Collagen I, Cat# 3447-020-01) and loaded into the ECM channel located in the middle lane. The plate was then incubated overnight at 37 °C and 5% CO₂. The following day, cells with or without 3-week pre-treatment of dTAG^V^-1 at concentrations ranging from 0 to 150 nM were detached using accutase, counted, and seeded at 5,000 cells/µL in 2 µL of complete DMEM medium into one of the perfusion channels adjacent to the ECM channel. This resulted in passive pumping as described previously (80), for a final concentration of 10,000 cells per chip. After 2 hours of cell attachment, 50 µL of complete medium containing the corresponding dTAG^V^-1 ligand concentration was added to the inlets and outlets of both perfusion channels to achieve the desired gradient concentrations. The plate was incubated at 37 °C and 5% CO₂ on a rocking platform (8-minute intervals, 14° inclination) for 7 to 10 days. The channels are separated by two phase guides (100 × 55 µm, *w* × *h*) that enables retention of ECM through pinning allowing cultured cells grown in a perfusion channel to be in direct contact with the ECM and invade through it in a barrier-free fashion (81). At the endpoint, the plates were fixed with 3.7% formaldehyde in HBSS for 15 minutes. Cells were then stained with Hoechst 3342 (Invitrogen, Cat# H3570) for nuclear visualization and ActinRed 555 (Invitrogen, Cat# R3711) to stain actin cytoskeleton. Images were acquired using the ImageXpress Confocal HT.ai (Molecular Devices) with either a 10× or 20× objective and Molecular Devices CellReporterXpress Automated Image Acquisition and Analysis Software (RRID:SCR_025681). The images were analyzed using custom Fiji plugins developed by Mimetas. The positions of the nuclei along the Y-axis of the chip were plotted to facilitate statistical analyses of the extent of invasion.

### Soft agar colony formation assays

Cells were seeded in 6-well plates with or without a 3-week pre-treatment using dTAG^V^-1 ligand at concentrations ranging from 0 to 150 nM. Following detachment with accutase, 15,000 cells/mL were suspended in complete growth medium and mixed with 0.7% agarose in a 1:1 ratio (Sigma-Aldrich, Cat# A9414). The cell-agarose mixture was supplemented with dTAG^V^-1 at the indicated final concentrations and then plated onto 12-well plates pre-coated with a bottom layer of 0.7% agarose. Cells were incubated at 37°C in a humidified atmosphere with 5% CO_2_ for 3 weeks with medium replenished every 3-4 days to maintain dTAG^V^-1 ligand levels. At the end of the incubation period, colonies were fixed and stained with 0.01% crystal violet in 4% formaldehyde/PBS for 2 hours. Excess stain was removed by a single wash with DPBS. Colony images were captured using a flatbed scanner, and quantification was performed by manually counting the total number of colonies per well.

### RNA-sequencing

Total RNA was extracted using the RNeasy Mini Kit (Qiagen, Cat# 74106), following the manufacturer’s instructions, including on-column DNase digestion. Quality control was performed using a DeNovix DS-11 FX spectrophotometer. Library preparation and sequencing were conducted by the Biomedical Sequencing Facility (BSF) at CeMM Research Center for Molecular Medicine of the Austrian Academy of Sciences. Libraries were prepared using the QuantSeq 3’mRNA-Seq V2 Library Prep Kit FWD for Illumina (Lexogen, Cat# 113.96) and sequencing was performed on an Illumina NovaSeq 6000 Sequencing System (RRID:SCR_016387), generating 100 bp single-end reads.

### Analysis of RNA-seq data

RNA-seq reads were processed using the nf-core/rnaseq pipeline (82) (version 3.14.0) with the GRCh38.p14 primary assembly as reference and GENCODE (RRID:SCR_014966) v46 annotation. In the pipeline we used the following extra arguments for QuantSeq: STAR (RRID:SCR_004463) aligner options “--alignIntronMax 1000000 –alignIntronMin 20 --alignMatesGapMax 1000000 -- alignSJoverhangMin 8 --outFilterMismatchNmax 999 --outFilterMultimapNmax 20 --outFilterType BySJout --outFilterMismatchNoverLmax 0.1 --clip3pAdapterSeq AAAAAAAA”, and Salmon option “--noLengthCorrection”. All data analysis was performed in R (version 4.3.1)(83). For differential expression analysis we used DESeq2 (RRID:SCR_015687) (version 1.42.1, Wald test) (84). We applied adaptive shrinkage to the reported fold changes using the ‘lfcShrink’ command with type ‘ashr’ (85). We set the cutoffs for differential expression to FDR < 0.05 and absolute log_2_FoldChange > 2. To convert mapped read counts to variance-stabilized gene expression, we used the DESeq2 function ‘vst’. For principal component analysis and visualization purposes, we averaged the expression of replicates and rescaled this expression per gene to range from 0 to 1. Principal component analysis for the one-day-treated samples was performed using DEGs only. For the long-term-treated samples we used all genes with standard deviation > 0.5. Gene set enrichment analysis was performed using hypergeometric tests as implemented in the package hypeR (version 2.0.0)(86). The set of DEGs was compared against a background of all genes that could be tested (those that DESeq2 could report an adjusted *p*-value for). The gene sets tested were KEGG (RRID:SCR_012773) 2021 (87) (only when comparing data from single-cell clones to parental bulk) and MSigDB Hallmark 2020 (88) as downloaded from (84–88) https://maayanlab.cloud/Enrichr/#libraries on 10/9/2023(89). For visualizations, we filtered the results to gene sets that had an FDR < 0.001 and log odds ratio > 3 at any shown comparison. To avoid enrichment odds ratios of infinity we imposed a weakly informative prior based on the Cauchy distribution (90).

### SLAM-sequencing

Approximately 4.5 million cells were seeded onto 10 cm dishes and grown to 80% confluency. The next day, cells were pre-treated with dTAG^V^-1 ligand at concentrations ranging from 0 to 150 nM for 3 hours to establish an EF protein gradient. Subsequently, newly synthesized RNA was labeled with 1 mM 4-thiouridine (4SU, Biosynth, Cat# NT06186) for 100 minutes and cells were harvested by snap-freezing plates on dry ice. RNA extraction was performed using the RNeasy Plus Mini Kit (Qiagen, Cat# 74134) according to the manufacturer’s instructions. Total RNA was subjected to alkylation with 10 mM iodoacetamide (IAA, Sigma-Aldrich, Cat# I6125) for 15 minutes and RNA was re-purified by ethanol precipitation. The purified RNA was quantified using a Qubit RNA broad range assay (Invitrogen, Cat# Q10210). 400 ng of alkylated RNA were used as input for generating 3’-end mRNA sequencing libraries using the QuantSeq 3’mRNA-Seq Library Prep Kit FWD for Illumina (Lexogen, Cat# 113.96) and PCR Add-on Kit for Illumina (Lexogen, Cat# 020). The distribution of RNA fragments was analyzed using High Sensitivity D1000 assay (Agilent, ScreenTape Cat# 5067-5548 and Reagents Cat# 5067-5585) in Agilent 4200 Tapestation. Libraries were pooled at 4 nM, and shallow sequencing was performed using Illumina MiSeq System (RRID:SCR_016379) to estimate the required read counts per sample, aiming for conditions where more than 3% of reads contained ≥2 T>C conversions. Deep sequencing was performed on an Illumina NovaSeq 6000 Sequencing System, generating 150 bp single-end reads.

### Analysis of SLAM-seq data

Where not stated otherwise, processing and analysis of the SLAM-seq data was performed as for the RNA-seq data. SLAM-seq reads were processed using the nf-core/slamseq pipeline (version 1.0.0) (91). The minimum number of T>C conversions in a read to call it a T>C read was set to 2. We used the count matrix of T>C reads (newly synthesized RNA) on the transcript level for downstream analysis. Differential expression of samples with various dTAG^V^-1 treatments compared to 0nM dTAG^V^-1 control was tested. Results were on the level of transcripts and to avoid multiple entries per gene, we kept the transcript with the lowest log *p*-value after averaging log *p*-values across all tests. We set the cutoffs for differential expression to FDR < 0.05 and absolute log_2_FoldChange > 1. Clustering of differentially expressed genes was performed on vst-normalized and rescaled data (see RNA-seq methods for details). We used the kmeans method (92) with 25 random starts for clustering. To determine the optimal number of clusters K, we varied K from 2 to 16 and calculated the gap statistic and followed the recommendations by CITE (93). GSEA for clusters was performed against a background of all genes that could be tested for differential expression at any given dTAG concentration.

### CUT&RUN

Cells were expanded into 10 cm dishes and pre-treated with dTAG^V^-1 ligand at concentrations ranging from 0 to 150 nM for 3 hours. Following treatment, cells were detached using accutase and subsequently frozen in aliquots of 650,000 cells per condition. The frozen cells were then subjected to the CUT&RUN protocol according to the manufacturer’s instructions using the CUTANA CUT&RUN Kit (EpiCypher, Cat# 14-1048). H3K4me3 and IgG antibodies served as positive and negative controls, respectively. A K-MetStat panel was added to these samples for quality control. E. coli DNA (0.25 ng) was used as spike-in control for library normalization. After purification, CUT&RUN DNA was quantified using a Qubit 1X dsDNA high sensitivity assay (Invitrogen, Cat# Q33230). A maximum of 5 ng DNA served as input for library preparation using CUTANA CUT&RUN Library Prep Kits with primer set 1 and 2 for multiplexing (EpiCypher, Cat# 14-1001 and Cat# 14-002). Of note, to avoid denaturing small fragments, end repair inactivation was performed at 50°C for 1 hour instead of 65°C for 30 minutes (94). The distribution of DNA fragment sizes was analyzed using High Sensitivity D1000 assay (Agilent, ScreenTape Cat# 5067-5548 and Reagents Cat# 5067-5585) in Agilent 4200 TapeStation System (RRID:SCR_018435). The molarity of each library was calculated and normalized to 5 nM. Libraries were pooled and sequenced using the Illumina NovaSeq 6000 Sequencing System, generating 50-bp paired-end reads.

### Analysis of CUT&RUN data

CUT&RUN reads were processed using the nf-core/cutandrun pipeline (version 3.2.2) with the GRCh38.p14 primary assembly as reference and Gencode v46 annotation. In the pipeline, we set the peak caller to MACS2 (95) with the broad peak setting for histone marks (H3K4me3, H3K27me3, H3K27ac) and narrow peak setting for all other samples. As a quality measure, we removed all peaks that overlapped regions that are known to cause high unspecific signal (96) (ENCODE4 exclusion list ENCFF356LFX). To remove spurious signal, we defined consensus peak regions per antibody target as follows: keep only significant peaks (FDR < 0.01) in every replicate, create union of peak ranges, keep only ranges that overlap significant peaks from at least two replicates. For FLI1 we combined the genotype-specific consensus peak ranges (parental and A2.2 clone) into one consensus range set by taking the union. To quantify the CUT&RUN signal at consensus peaks, we counted the fragments that map to those regions using the bamsignals R package (v 1.34.0). In subsequent analysis steps that require quantitative comparisons of conditions, the resulting peak read count matrix is analyzed with the same tools that we used for RNA-seq. To annotate the consensus peak regions with respect to their genomic location, we used the ChIPseeker (97) R package version 1.39.0 with the promoter region defined as 1kbp up- and downstream of the TSS, and the gene flanking region length set to 5kbp. We further annotated the consensus peaks with transcription factor binding motifs (TFBS) occurrence information. To do so, we used the motifmatchr R package version 1.24 to scan for all matches of the JASPAR2020 (45) human core motifs (matchMotifs function with default p-value cutoff of 5e-05). To visualize the heterogeneity of the FLI1 consensus peaks based on the presence of TFBS, we transformed the high-dimensional data (12,370 peaks by 633 motifs) to a two-dimensional space. For this we used UMAP(98) with the matrix of match counts as input and the following parameters: metric = “manhattan”, n_neighbors = 100, ret_nn = TRUE, pca = 10, pca_center = TRUE. To group the peaks, we calculated the shared nearest neighbor graph (Manhattan distance, 100 neighbors), pruned edges smaller than 1/15, and applied the community detection Louvain clustering (99). For these steps we used implementations in the Seurat R package version 4.3.0.1 (100). To group the peaks, we calculated the shared nearest neighbor graph (Manhattan distance, 100 neighbors), pruned edges smaller than 1/15, and applied the community detection Louvain clustering (99). For these steps we used implementations in the SEURAT (RRID:SCR_007322) R package version 4.3.0.1(100). We explored potential links of peak clusters with SLAM-seq gene clusters using enrichment tests. Specifically, we assigned every FLI1 consensus peak to its closest expressed gene (TSS within 150kbp) and, if the gene was in one of the five SLAM-seq clusters, assigned the gene cluster to the peak. This way we assigned 1,128 peaks to SLAM-seq clusters. Using this set of peaks, we then tested all peak cluster and gene cluster combinations for association (two-sided Fisher’s exact test).

### Animal experiments and ethics

Mice were bred and housed under pathogen-free conditions at a temperature of 22 ± 1°C, with 55 ± 5% humidity, and a 14-hour light/10-hour dark photoperiod. *NOD.Cg-Prkdc^scid^ Il2rg^tm1Wjl^/SzJ* (NSG, RRID: BCBC_4142) mice were obtained from the in-house breeding facility of the Vienna Biocenter. All mouse experiments were conducted in accordance with a license approved by the Austrian Ministry (GZ: MA58-2260492-2022-22). Transparent larvae from Casper zebrafish (*Danio rerio*) background (*mitfa*^w2/w2^; *mpv*^a9/a9^) stably expressing mCherry to label blood vessels (*Tg(kdrl:Hsa.HRAS-mCherry*)^s896^) were generated from parental fish pairs bred and maintained under standard conditions at the CCRI zebrafish facility (101), in accordance with the license GZ:565304-2014-6 of the local authorities.

### In vivo metastasis assay

Luciferase-expressing A2.2 EF-dTAG cells were pretreated with or without 15 nM dTAG^V^-1 ligand for 3 weeks. An additional condition was included, where cells pretreated with 15 nM dTAG^V^-1 for 3 weeks underwent a 7-day washout period. Following treatment, cells were detached using Accutase, resuspended in sterile PBS containing a final concentration of 15 nM dTAG^V^-1 (for the pre-treated condition), and injected intravenously via the tail vein into male NSG mice aged 7-13 weeks. A total of 1 million cells in a final volume of 100 μL were injected per mouse. Five mice were injected for each of the following conditions: A2.2 without dTAG^V^-1 (Control), A2.2 with 15 nM dTAG^V^-1, and A2.2 with 15 nM dTAG^V^-1 followed by a 7-day washout (EF rescue), as well as parental A673 cells cultured for the same duration. Mice were imaged weekly using the IVIS Spectrum Xenogen bioluminescent imaging system (Caliper Life Sciences) and no further dTAG^V^-1 ligand was supplied to the mice during the entire duration. Mice were anesthetized with 2-3% isoflurane inhalation and injected retro-orbitally with D-luciferin (150 mg/kg; Gold Biotechnology). Bioluminescent images were captured 2 minutes post-injection to assess metastatic burden based on the bioluminescent signal (BLI). Mice were euthanized upon reaching humane endpoints (e.g., weight loss > 20%, signs of distress or pain) or when the BLI signal increased by at least 100-fold compared to day 0. At the experimental endpoint, animals were anesthetized, injected with D-luciferin, and then euthanized. Lungs and livers were imaged using the IVIS system before being collected for formalin-fixed paraffin-embedded (FFPE) processing and subsequent Hematoxylin and Eosin (H&E) staining. Imaging data were analyzed using Living Image software (RRID:SCR_014247) (v.4.8).

### Zebrafish experiments

A2.2 EF-dTAG cells were pretreated with either 5 nM or 15 nM dTAG^V^-1 ligand for 3 weeks. An additional condition involved cells that were pretreated with 15 nM dTAG^V^-1 for 3 weeks and subsequently underwent a 7-day washout period. Following treatment, cells were detached using Accutase and stained with CellTrace™ Violet (Invitrogen, Cat# C34571) for visualization. Two-day post-fertilization (dpf) zebrafish larvae were anesthetized using 1x tricaine (pH 7.2; 0.16 g/L Ethyl-3-aminobenzoate methanesulfonate (Sigma Aldrich, Cat# E1052110G) in E3 medium). Cells from the four different conditions (untreated, 5 nM dTAG^V^-1, 15 nM dTAG^V^-1, and 15 nM dTAG^V^-1 with washout) were resuspended in PBS, with or without dTAG^V^-1 ligand, depending on the treatment or washout conditions. To facilitate observation of the extravasation process from the tail vessels, 200-300 cells were injected into the duct of Cuvier using a Eppendorf FemtoJet 4i (RRID:SCR_019870) to directly access the circulation, as previously described (102). Twenty-four xeno-transplanted larvae per condition were sorted for a roughly similar cell load in circulation 2 hours after injection, as observed under a fluorescence microscope. The larvae were then incubated at 34°C without further addition of the dTAG^V^-1 ligand to the water. Larvae were aligned in a 96-well plate (ZFA 101-02a, Funakoshi) and imaged 1- and 3-days post-injection using the Operetta CLS high-content imager (RRID:SCR_018810, Revvity, formerly PerkinElmer). Images of the tail region were acquired with a 5x objective in confocal mode and evaluated using Harmony Software 4.9. This analysis located red and blue fluorescence above set threshold to delineate the blood vessel area and define the cell area, allowing for the generation of an overlapping mask to determine the number of cells located outside or inside the vessel area.

### Survival analysis

Kaplan-Meier survival analyses were carried out in 196 molecularly confirmed EwS patients and retrospectively collected primary tumors were profiled at the mRNA level by gene expression microarrays in previous studies ^46-49^. To this end, microarray data generated on Affymetrix HG-U133Plus2.0, Affymetrix HuEx-1.0-st or Amersham/GE Healthcare CodeLink microarrays of the 196 EwS tumors (Gene Expression Omnibus (GEO) accession codes: GSE63157 ^46^, GSE12102 ^47^, GSE17618 ^48^, GSE34620 ^49^ provided with clinical annotations were normalized separately as previously described ^102^. Only genes that were represented on all microarray platforms were kept for further analysis. Batch effects were removed using the ComBat (RRID:SCR_010974) algorithm^103^. Data processing was done in R (Version 3.4.2).

### Data analysis, statistics and reproducibility

All experiments were performed in at least two independent biological replicates unless stated otherwise. The sample size (*n*) refers to biological replicates. Data are presented as the mean when *n* = 2 and as the mean ± standard error of the mean (SEM) when *n* ≥ 3. Statistical analyses were conducted using one-way analysis of variance (ANOVA), two-way ANOVA with Dunnett’s post hoc multiple comparisons test, as described in the figure legend for each experiment. For mouse survival analyses, significance was determined using the log-rank (Mantel–Cox) test. Statistical analyses were performed using GraphPad Prism (version 9.41, GraphPad Software) and R (version 4.3.1)(83). Survival analyses were conducted using the Kaplan–Meier method. Data were assumed to follow a normal distribution, except for transwell migration and soft agar assays, which were log10 or log10+1 transformed prior to statistical testing. Graphs were generated using GraphPad Prism (RRID:SCR_002798) (version 9.41) and R (version 4.3.1). Statistical significance was defined as *P < 0.05*.

## Data and materials availability

The data generated in this study have been deposited in Gene Expression Omnibus (GEO) (RRID:SCR_005012) (https://www.ncbi.nlm.nih.gov/geo/query/acc.cgi?acc=GSE291136) at GSE291136.

## Acknowledgements

We are grateful to Georg Winter and Dave Aryee for providing constructs for cloning of the donor plasmids, and Karin Mühlbacher for technical assistance. We thank Johannes Zuber for helping with SLAM-seq optimization. We also acknowledge the CCRI FACS core facility for support with cell sorting and the ZANDR platform for assistance with zebrafish extravasation experiments.

## Author contributions

Conceptualization, H.K. Data curation, V.S., V.F., C.H., A.K. and S.G., Formal Analysis, C.H., V.S., C.S. F.H. and F.C.-A. Funding acquisition, H.K., T.G., F.H. and A.O. Investigation, V.S., V.F., A.K., S.G., A.W.-W., F.C.-A. and M.M.Z. Methodology, V.S., V.F., K.Q., and D.K. Project administration, H.K. Resources, F.H., A.O., M.D., T.G., J.A., and A.S. Supervision, H.K., V.F., F.H., M.D., A.O., T.G. and K.Q. Visualization, V.S., V.F. and C.H. Writing – original draft, V.F., V.S., C.H., and H.K. Writing – review & editing, H.K., F.H., V.S., V.F., C.H., A.K., C.S., F.C.-A., T.G., M.D. and A.O

