## Supplementary Figures for "Dynamic modelling of EWS::FLI1 fluctuations reveals molecular determinants of phenotypic tumor plasticity and prognosis in Ewing sarcoma"

**Supplementary Figure S1.**

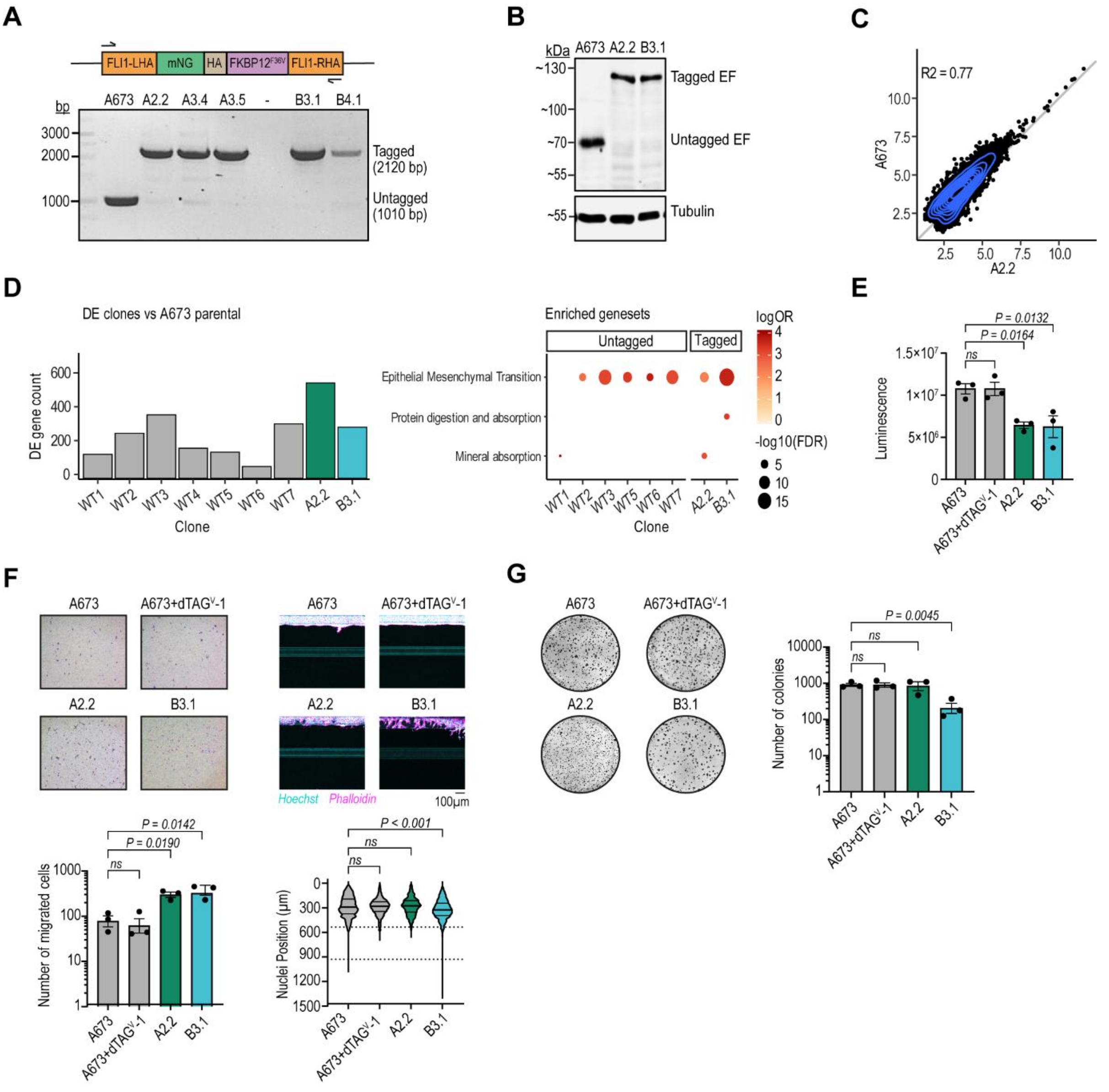


Characterization of EF-dTAG clones. **A**, Schematic depicting primer locations (arrows) on the FLI1-LHA-mNG-HA-FKBP12$^{F36V}$–FLI1-RHA homology-directed repair template used for clonal genotyping of the FLI1 locus (top). Representative agarose gel images showing products of two-step RT-PCR, confirming the presence of the tagged *EWSR1::FLI1* fusion transcript in five EF-dTAG clones with bi-allelic knock-in, using the indicated primers (bottom). **B**, Western blot analysis of untagged and tagged EF protein levels in parental A673 cells and EF-dTAG clones. Wildtype FLI1 (~52 kDa) was undetectable in all samples tested. Tubulin was used as a loading control. Data shown are representative of at least three independent experiments. **C**, FLI1 CUT&RUN-seq peak signal in A2.2 clone and

A673 parental line. Each dot is one of 12,370 consensus peaks. Values indicate normalized read counts, averaged across replicates. **D**, Number of differentially expressed genes (DEGs) between single-cell clones and A673 parental line (left, DESeq2[74], FDR < 0.05, absolute $\log_2$ fold change > 1). Functional enrichment of DEGs identifies MSigDB Hallmark 2020 and KEGG 2021 pathways as significantly enriched at least once (right, hypergeometric test, FDR < 0.001, $\log_2$ odds ratio > 3, background: all DESeq2 tested genes). **E**, Proliferation of EF-dTAG clones and parental Aä673 cells measured using a CellTiter-Glo assay. Data are presented as mean ± SEM (*n* = 3 independent experiments). **F**, Transwell migration assays showing migrated cells stained with crystal violet 24 hours after seeding (top left) and organoplate invasion assays depicting cell invasion into collagen I extracellular matrix (ECM) 7 days after seeding EF-dTAG clones and parental A673. Cells were pre-treated with 150nM $dTAG^{V}$-1 for 24 hours where indicated. Cells were stained with Hoechst (cyan, nuclei) and phalloidin (magenta, actin cytoskeleton) (top right). Representative images from one of three independent experiments are shown, scale bars, 100 µm. Migrated cell counts are presented as mean ± SEM after log10 transformation (bottom left; *n* = 3). Nuclei positions along the Y-axis in the organoplate invasion assays are depicted, with phase guides shown as dotted lines at distances of 535 µm and 930 µm (bottom right; *n* = 3). **G**, Soft agar colony formation assays showing crystal violet stained colonies formed by EF-dTAG clones and parental Aä673 cells three weeks after seeding. Representative images from one of three independent experiments are shown. Colony counts are presented as mean ± SEM (*n* = 3) following log10+1 transformation. For panels (**E-G**), *P*-values were calculated using one-way ANOVA with Dunnett's post-hoc multiple comparisons test. *ns*, not significant.

**Supplementary Figure S2.**

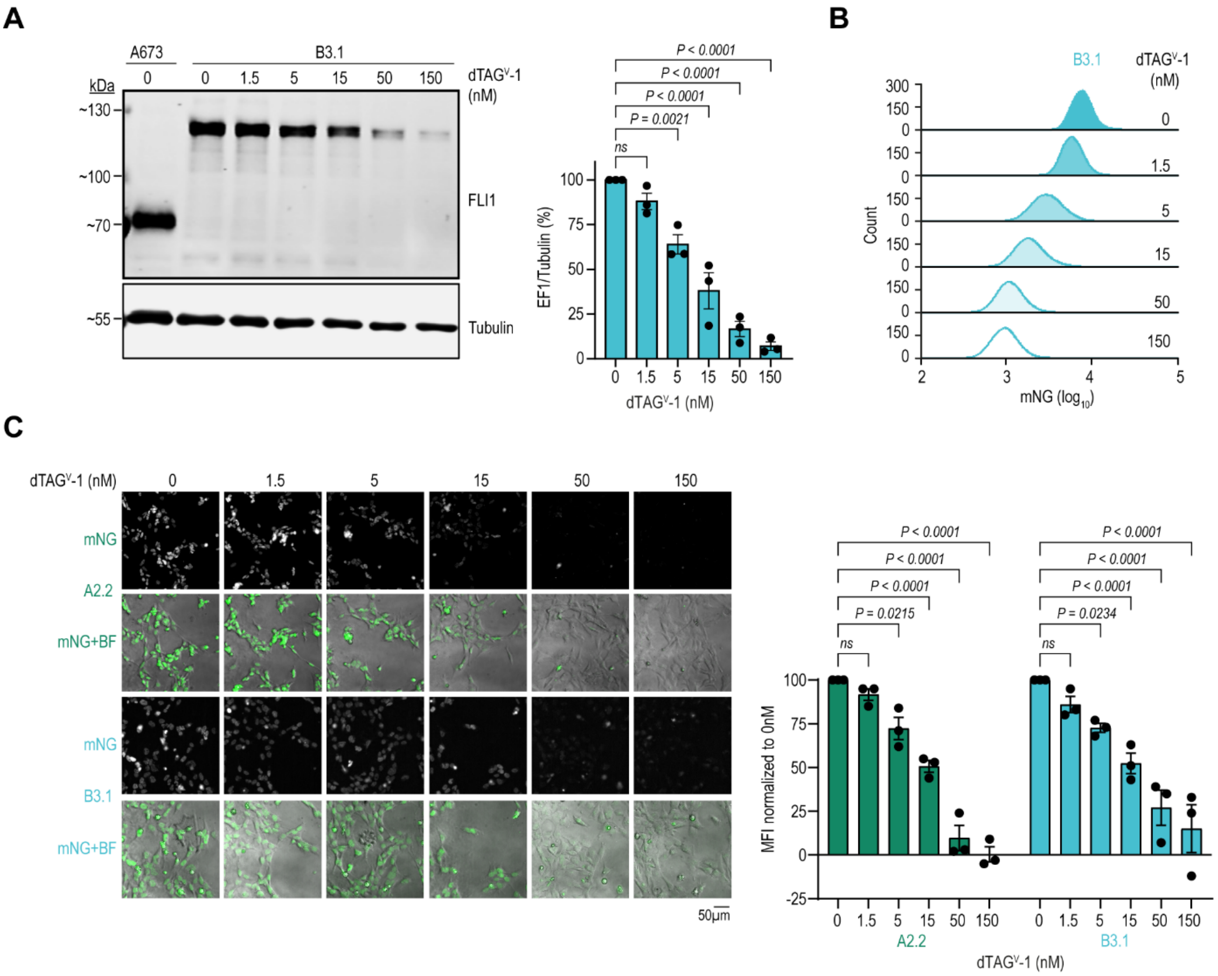


EF gradient characterization of EF-dTAG clones. **A**, Western blot analysis of FLI1 protein levels in EF-dTAG clone B3.1 and parental A673 cells treated with the indicated concentrations of dTAG$^{V}$-1 for 24 hours. Tubulin was used as a loading control. Bar graphs depict EF protein levels normalized to Tubulin, presented as mean ± SEM (right; *n* = 3 independent experiments). *P*-values were calculated using One-way ANOVA with Dunnett's post-hoc multiple comparisons **B**, Flow cytometry analysis of mNG fluorescence intensity in EF-dTAG clone B3.1 following treatment with increasing concentrations of dTAG$^{V}$-1 concentrations for 24 hours, data are representative of at least three independent experiments. **C**, Confocal images for mNG fluorescence in EF-dTAG clone A2.2 (top) and B3.1 (bottom) treated with the indicated concentrations of dTAG$^{V}$-1 for 24 hours, scale bars, 50 μm. Images are representative of at least three individual experiments; BF, Brightfield. Bar graphs represent mean mNG mean fluorescence intensity values normalized to the 0 nM control, presented as mean ± SEM (right; *n* = 3 independent experiments). *P*-values were calculated using Two-way ANOVA with Dunnett’s post-hoc multiple comparisons test. *ns*, not significant.

**Supplementary Figure S3.**

**A**

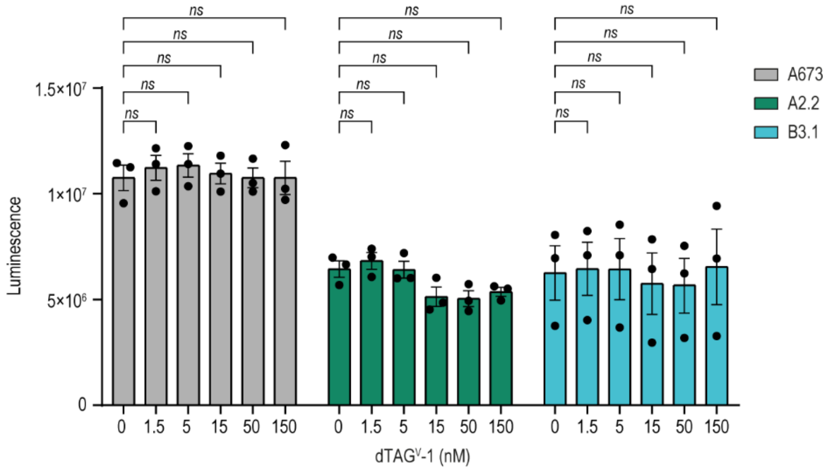


**B**

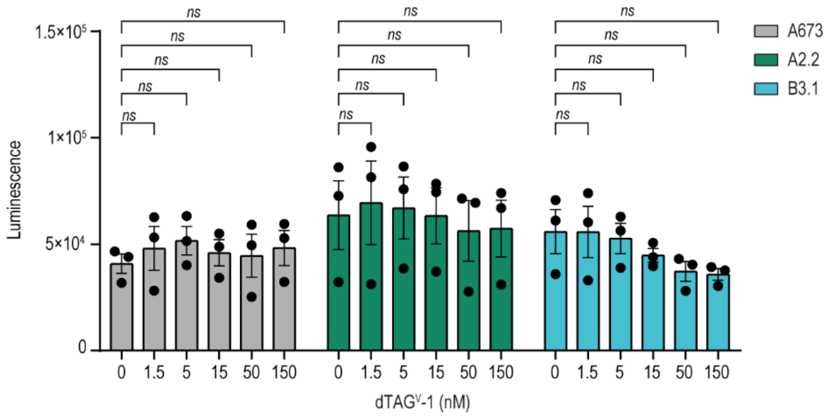


Effects of dTAG$^{V}$-1 on proliferation and apoptosis. **A**, Proliferation of EF-dTAG clones and parental A673 cells in response to increasing concentrations of dTAG$^{V}$-1, assessed using the CellTiter-Glo assay. Data represents the mean ± SEM of *n* = 3 independent experiments. **B**, Apoptosis of EF-dTAG clones and parental A673 cells in response to increasing concentrations of dTAG$^{V}$-1, measured using the Caspase-Glo 3/7 assay. Data represents the mean ± SEM of *n* = 3 independent experiments. *P*-values for both panels were determined using two-way ANOVA with Dunnett's multiple comparisons test. *ns*, not significant.

**Supplementary Figure S4.**

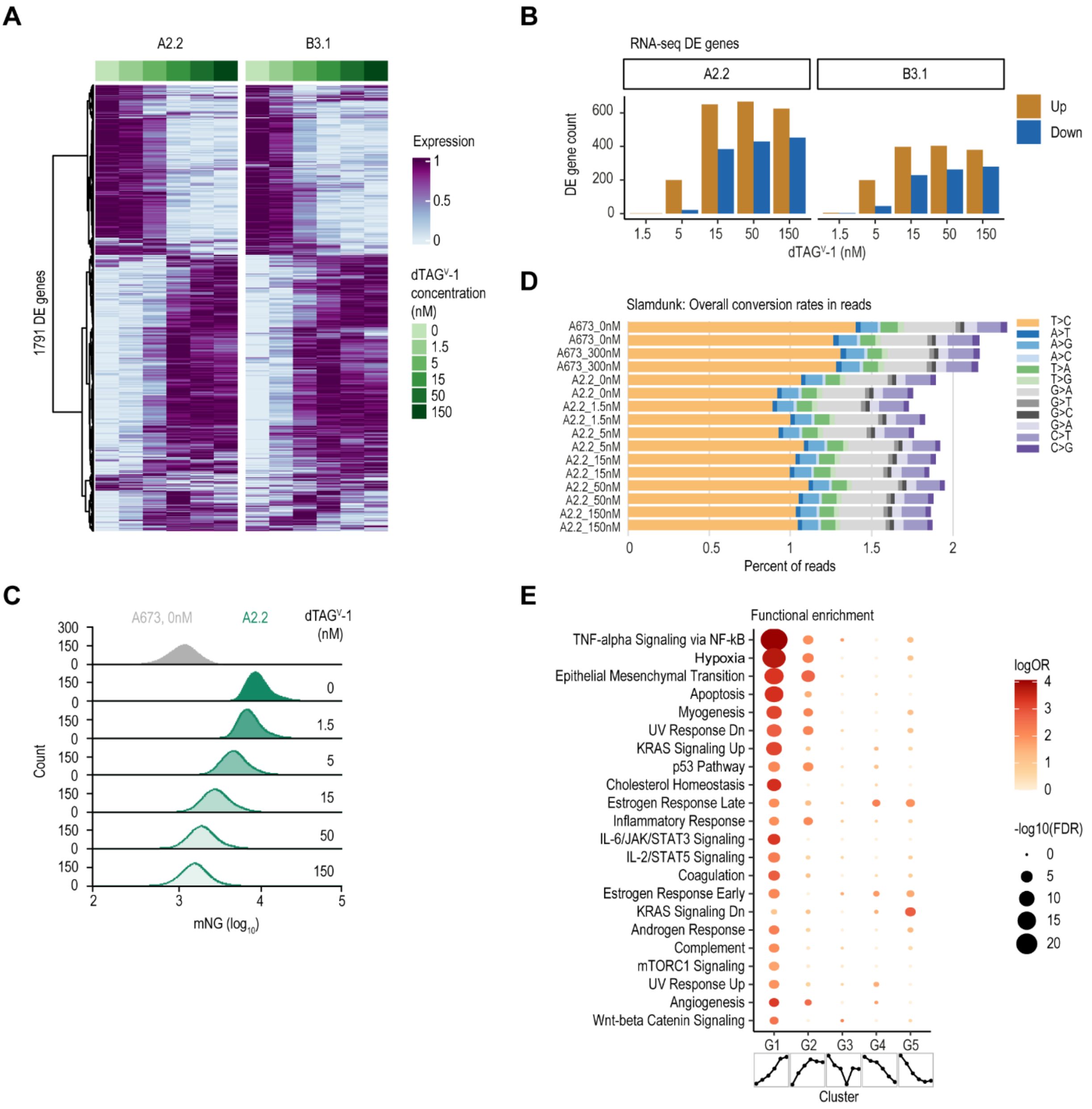


Transcription profiling of EF-dTAG clones in response to dTAG$^{V}$-1 treatment. **A**, Heatmaps showing gene expression changes in the 24-hour dTAG$^{V}$-1 gradient treatment experiment. Y-axis shows all 1791 genes determined differentially expressed (DESeq2[74], FDR < 0.05, absolute $\log_2$ fold change > 2) in at least one EF-dTAG clone at any dTAG$^{V}$-1 concentration. Gene expression was normalized using DESeq2 vst, averaged across replicates, and rescaled per gene per clone. **B**, Number of DEGs from panel (**A**) across treatment conditions. **C**, Flow cytometry analysis of mNG fluorescence intensity in EF-dTAG clone A2.2 after 3 hours of treatment with increasing dTAG$^{V}$-1 concentrations, representative of samples sequenced in SLAM-seq. **D**, SLAM-seq base conversion rates in percent of all reads as reported by the Slamdunk processing pipeline. **E**, Functional enrichment analysis of SLAM-seq cluster genes, showing MSigDB Hallmark 2020 pathways that were significantly enriched (hypergeometric test, FDR < 0.05, $log_2$ odds ratio > 1, background: all detected genes) in at least one cluster. Icons below the cluster labels show the mean expression pattern per cluster (x-axis shows increasing dTAG$^V$-1 concentrations, y-axis rescaled expression).

**Supplementary Figure S5.**

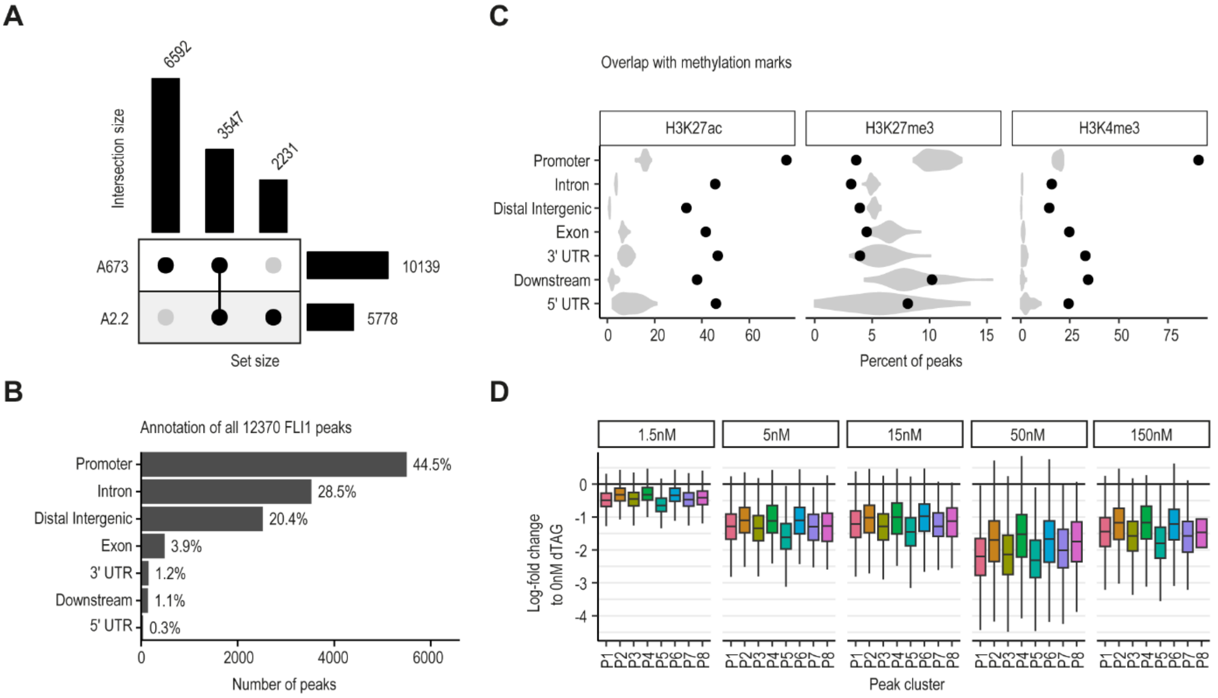


Characterization of FLI1 binding peaks and dynamics of peak signals. **A**, UpSet plot showing the intersection of the CUT&RUN FLI1 peaks discovered in A2.2 and parental A673 cells. **B**, Annotation of FLI1 peaks with respect to genomic location. **C**, Percentage of FLI1 peaks (x-axis), grouped by genomic location (y-axis), overlapping methylation marks. Observed percentages are shown as black points and random background distributions of shuffled peaks (all peaks randomly placed in genome 50 times) depicted as gray violin plots. **D**, FLI1 CUT&RUN peak signal change for all FLI1 consensus peaks grouped by peak cluster and $dTAG^{V}$-1 concentration in A2.2 clone. Boxplots show log-fold change compared to 0nM condition as reported by DESeq2[74] with ashr[76] shrinkage applied.

**Supplementary Figure S6.**

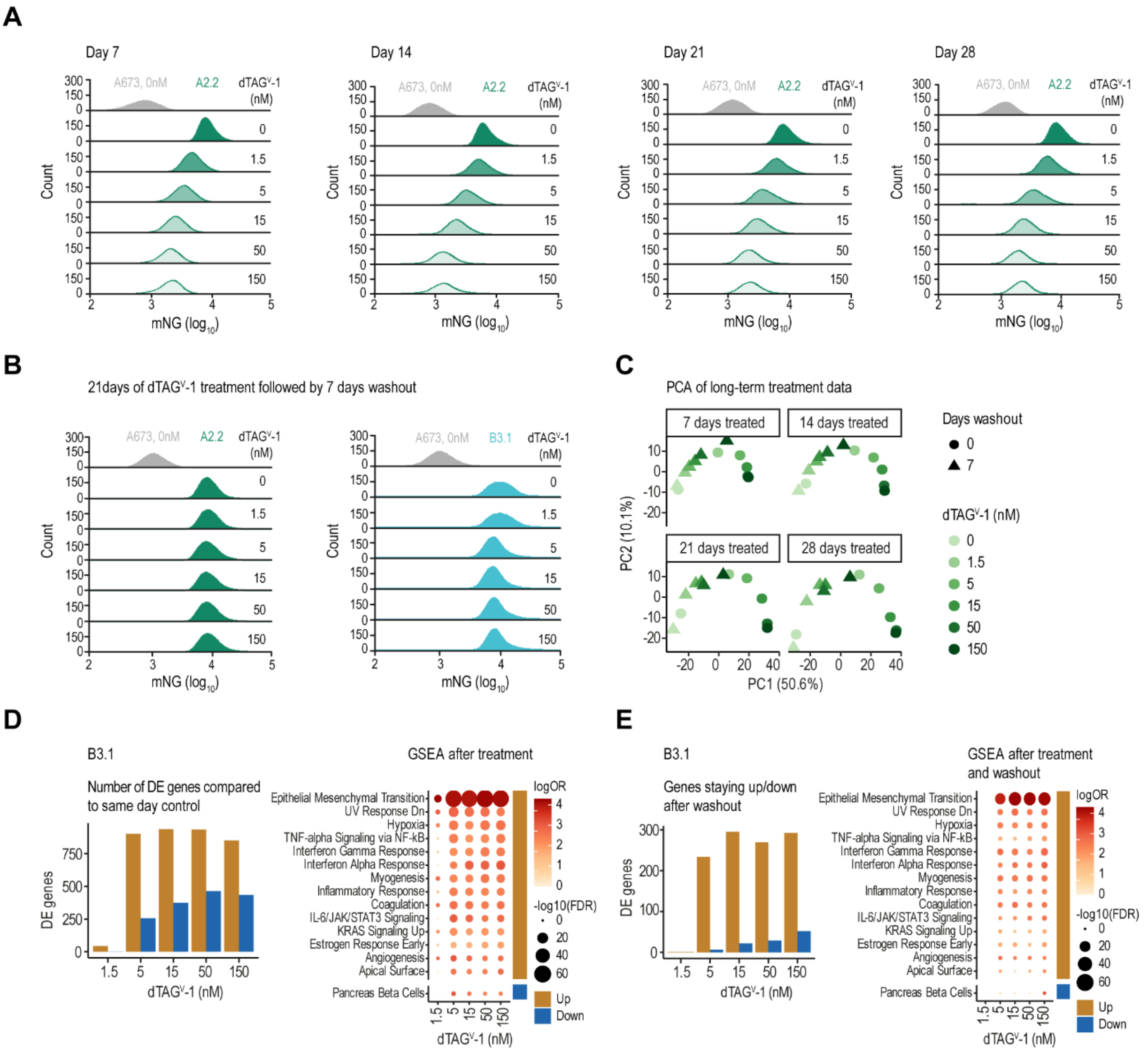


Long term EF gradient treatment and transient effects due to dTAG$^{V}$-1 treatment. **A**, Flow cytometry analysis of mNG fluorescence intensity in EF-dTAG clone A2.2 after 7, 14, 21, and 28 days of treatment with increasing concentrations of dTAG$^{V}$-1. Representative data corresponds to samples used for RNA-sequencing. **B**, Flow cytometry analysis of mNG fluorescence intensity in EF-dTAG clones after 21 days of dTAG$^{V}$-1 treatment at the indicated concentrations followed by a 7-day washout period. **C**, PCA of prolonged and transient EF dosage modulation on transcription in A2.2 clone (RNA-seq), showing first two principal components calculated using 1,455 highly variable genes (gene expression with standard deviation > 0.5 after normalization with DESeq2[74] vst function and averaging two replicates). **D**, Number of DEGs (Left, DESeq2, FDR < 0.05, absolute $log_2$ fold change > 2) and functional pathways enrichment analysis (Right) for clone B3.1 after 21-day treatment with increasing concentrations of dTAG$^{V}$-1. Pathways shown are

MSigDB Hallmark 2020 pathways that were significantly enriched (hypergeometric test, FDR < 0.001, $\log_2$ odds ratio > 3) in any treatment condition. **E**, As **D** but showing results for treatment followed by 7-day washout.

**Supplementary Figure S7.**

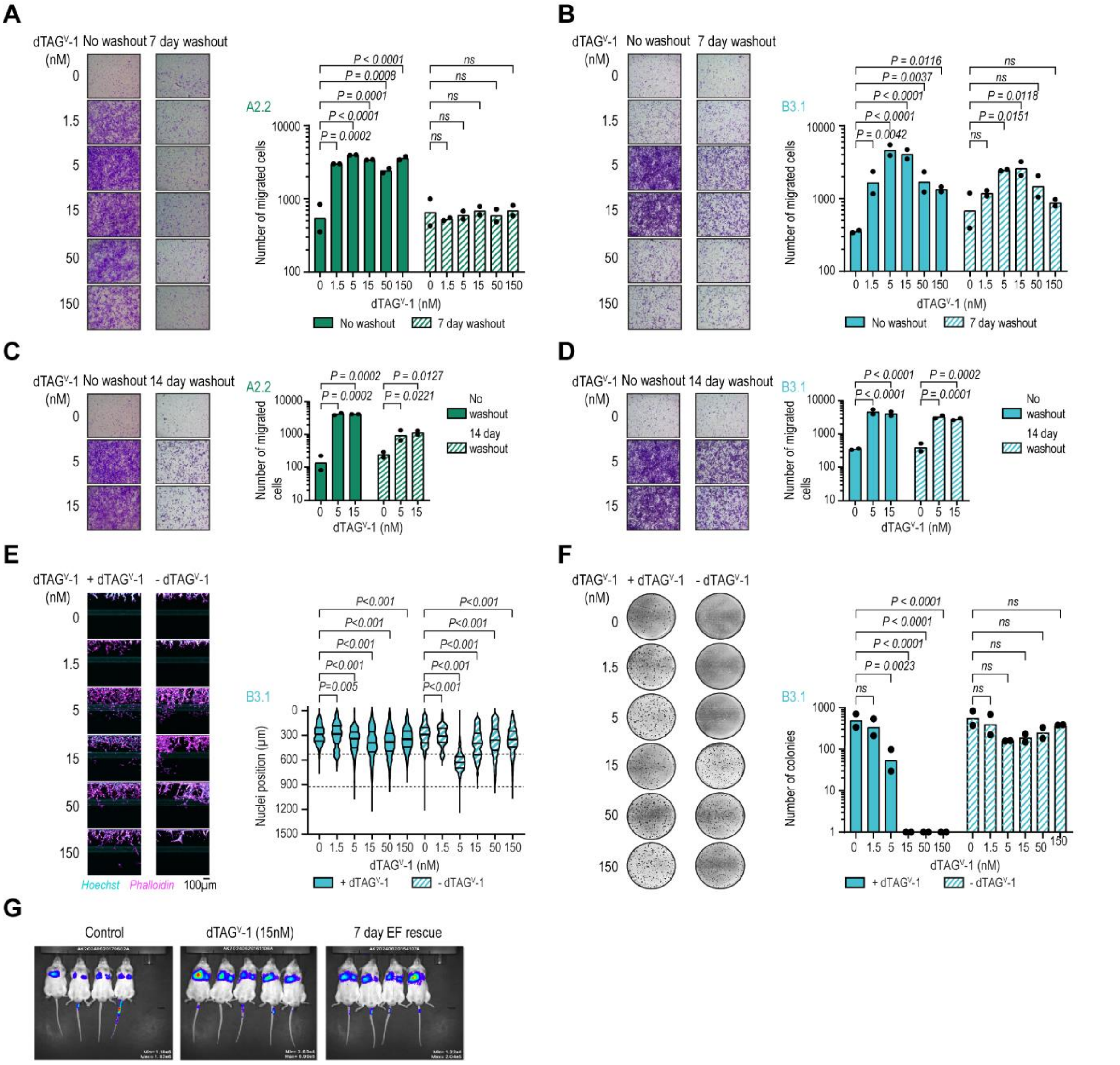


Metastatic phenotypic analysis of EF-dTAG clones following 21-day dTAG$^{V}$-1 treatment and washout. **A**, Transwell migration assays showing migrated A2.2 cells after 7-day treatment with dTAG$^{V}$-1 at the indicated concentrations or treatment for 7 days followed by a 7-day washout. Representative images from one of two independent experiments are shown (left). Migrated cell counts are presented as mean after log10 transformation (right; *n* = 2). **B**, Transwell migration assays showing migrated B3.1 cells following treatment with dTAG$^{V}$-1 for 21 days at the indicated concentrations, or treatment for 21 days followed by a 7-day washout. Representative images from one of two independent experiments are shown (left). Migrated cell counts, presented as mean after log10 transformation, are shown on the right (*n* = 2). **C**,**D** Transwell migration assays showing migrated A2.2 cells (**C**) or B3.1 cells (**D**) following 21-day treatment with dTAG$^{V}$-1 at the indicated concentrations or following 21 days of treatment and a subsequent 14-day washout. Representative images from one of two independent experiments are shown (left). Migrated cell counts are presented as mean after log10 transformation (right; $n = 2$). **E**, Organoplate invasion assays showing cell invasion into collagen I ECM 7 days after seeding B3.1 cells treated with dTAG$^{V}$-1 for 21 days at the indicated concentrations, with or without further dTAG$^{V}$-1 treatment. Representative images from one of two independent experiments are shown (left). Nuclei positions along the Y-axis are shown, with phase guides indicated by dotted lines at 535 µm and 930 µm (right; $n = 2$). **F**, Soft agar assays showing colonies formed by B3.1 cells three weeks after seeding 21 days dTAG$^{V}$-1 treated cells at indicated concentrations with or without further dTAG$^{V}$-1 treatment. Representative images from one of two independent experiments are shown (left). Colony counts are presented as mean ($n = 2$) following log10+1 transformation. **G**, Representative IVIS images of mice showing the homing of luciferase expressing A2.2 cells to lungs directly after (Day 0) tail vein injection of cells treated with 15 nM dTAG$^{V}$-1 for 21 days, with or without a 7-day washout along with control. *P*-values for all panels were calculated using two-way ANOVA with Dunnett's post-hoc multiple comparisons test. *ns*, not significant.
